# Spatial metabolic ecosystems shape the architecture of *IDH*-mutant glioma

**DOI:** 10.64898/2026.11.15.765028

**Authors:** Joel J.D. Moffet, Jurgen Kriel, Tianyao Lu, Montana Spiteri, Oluwaseun E. Fatunla, Adam L. Valkovic, Jasmine F. Pignatelli, Zachery Moore, Bhavika Kumar, Shana Portelli, Lucy A. Riley, Vinod K. Narayana, Brunda Nijagal, Jordan J. Jones, Sarah A. Cain, Samuel J. Roberts-Thompson, Mengbo Li, Katharine J. Drummond, Malcolm J. McConville, David M. Ashley, James R. Whittle, Saskia Freytag, Sarah A. Best

**Author notes:** These authors contributed equally (Joel J.D. Moffet, Jurgen Kriel, Tianyao Lu, Montana Spiteri). Senior author (James R. Whittle, Saskia Freytag, Sarah A. Best).

## Abstract

The principles governing tissue organization in *IDH*-mutant glioma remain poorly understood, particularly in the absence of hypoxia and necrosis that define *IDH*-wildtype glioblastoma. Here, we integrated spatial transcriptomics, metabolomics and histology across co-registered serial tissue sections, generating a multidimensional atlas of *IDH-*mutant glioma. Integration with an independent spatial transcriptomics cohort and single cell reference data, enabled resolution of metabolic and cellular heterogeneity underlying tissue organization. We identified twelve conserved spatial niches with distinct metabolic identities that could not be explained by cellular composition or histology alone. Beyond these ecosystems, we discovered a transcriptionally and epigenetically intermediate malignant cell state enriched in WHO grade 2 and 3 oligodendroglioma, and spatially segregated myeloid programs. Importantly, progression following chemoradiotherapy was associated with expansion of a glutathione-rich, NRF2-associated antioxidant niche. Together, our findings establish metabolic specialization as a fundamental dimension of glioma organization, providing new insight into tumor heterogeneity, progression and treatment-associated adaptation.

**HIGHLIGHTS:**

- Spatial co-registered multi-modal atlas maps niches in IDH-mutant glioma
- Niche identity is driven by metabolism, not by cellular composition
- A *RIT2*+ intermediate malignant state is enriched in oligodendroglioma
- A glutathione-rich antioxidant niche expands following treatment

## INTRODUCTION

Among adult gliomas, astrocytomas and oligodendrogliomas are the most common brain tumor in adults under 50^1–3^, and are defined by mutations in *IDH1/2* (mIDH) that drive accumulation of the oncometabolite D-2-hydroxyglutarate (2-HG)^4^, reshaping cellular differentiation, epigenetic state and tumor evolution^5^. Recent studies have shown that, similar to *IDH*-wildtype glioblastoma (GBM), these tumors comprise a set of malignant cell states, including astrocyte-like (AC-like), oligodendrocyte precursor-like (OPC-like), neural progenitor-like (NPC-like), mesenchymal-like (MES-like) and cycling progenitor populations, interwoven with cells of the brain microenvironment^6–8^. These cellular populations are spatially patterned by the underlying normal brain architecture, with white and gray matter structures shaping local cellular composition^9^. While normal brain anatomy provides an important scaffold for tumor growth, the association of specific malignant and immune cell states suggests that additional forces shape tissue organization^10–12^. In WHO grade 4 GBM, these factors include hypoxia and necrosis^12,13^, however it is unclear whether equivalent organizing principles exist in grade 2 and 3 mIDH gliomas^14^, as these lower-grade tumors develop in the absence of extensive hypoxia and necrosis^6,15^. Given cellular metabolism is fundamentally rewired in mIDH tumors, metabolic organization represents a compelling but largely unexplored organizing principle.

Understanding these relationships has become increasingly important with the introduction of mIDH inhibitors into clinical practice^16^. In a recent perioperative study, four-weeks of mIDH inhibition induced profound metabolic remodeling, altered cycling progenitor tumor cell programs, promoted differentiation, and increased immune infiltration, demonstrating that therapeutic response is closely coupled to metabolic state^17^. Beyond these acute treatment responses, gliomas undergo progressive molecular changes over years of disease evolution, particularly following chemoradiotherapy^18^. These include loss of CpG island methylation, acquisition of genetic alterations such as *PDGFRA* amplification, *CDKN2A* loss and treatment-associated hypermutation, and extensive remodeling of the immune microenvironment^6,7,15,19^. These events are associated with reduced differentiation and expansion of stem-like populations^15^. Remarkably, recurrence is accompanied by relatively limited changes in overall cell state composition, with recurring tumors retaining the same major malignant populations despite extensive epigenetic, genetic and microenvironmental remodeling^6,20^. This occurs irrespective of resistance to standard of care therapy^21,22^, suggesting that higher-order features of tumor organization, beyond cellular composition alone, play an important role in governing disease behavior and treatment response.

To address this gap, we constructed a multi-modal spatial atlas of grade 2 and 3 mIDH glioma by integrating spatial metabolomics (SM), spatial transcriptomics (ST) and immunofluorescence (IF) across serial tissue sections using a computational framework that enables high-resolution mapping of metabolites to specific cell populations and tissue niches^23,24^. We applied this approach to five deeply profiled tumors spanning up to eight aligned serial levels and extended our analyses through an additional ST cohort of 21 mIDH tumors, including longitudinal samples, anchored to a single-cell reference atlas comprising 101 samples. This integrated resource reveals distinct metabolic niches emerging following treatment, identifies an intermediate *RIT2+* malignant state bridging AC- and OPC-like populations, and defines changes in the myeloid microenvironment associated with disease progression. Together, these findings establish metabolism as a fundamental pillar of tumor organization in mIDH glioma.

## RESULTS

### Building a multi-dimensional spatial atlas of IDH-mutant glioma

Fresh tumor tissue from five WHO grade 2 and 3 mIDH astrocytomas were collected directly from the operating room and rapidly flash-frozen in an isopentane liquid nitrogen bath to preserve endogenous metabolites (**Table 1, Figure S1A**). To encompass the inherent intra- tumoral spatial heterogeneity, each tumor block was sampled across ∼480 µm using up to eight serial levels spaced 60 µm apart, with three adjacent 10 µm sections at each level (**Figure 1A**). Adjacent sections were profiled by histopathology, Xenium ST using a custom 307-gene panel (**Table S1**) at optical resolution followed by IF for GFAP, MAP2, actin and DAPI, and Shimadzu MALDI-TOF SM at 10 µm spatial resolution. Metabolite annotations were supported by orthogonal validation of twelve key metabolites using chemical standards, allowing comparison of their experimentally measured mass-to-charge (m/z) ratios and fragmentation patterns with those detected in patient tissue (**Figure 1B, S1B, Table S2**). Remaining tissue from the profiled tumor block or additional matched tissue from the same tumor underwent single-nucleus RNA sequencing (snRNA-seq). After annotation using marker genes (**Figure S1C, D**), matched ST and snRNA-seq showed concordance (mean correlation = 0.83, **Figure S1E**). Fine-scale alignment of adjacent multi-modal sections using SMINT enabled integration of cellular, histological and metabolic information across serial tissue sections (**Figure 1C**). We then combined ST profiles from the deeply profiled cohort with an additional 21 mIDH glioma samples from 10 patients (n = 15 patients total), comprising approximately 7 million nuclei in total (**Table 1, Figure S1A, F**).

**Figure 1.**
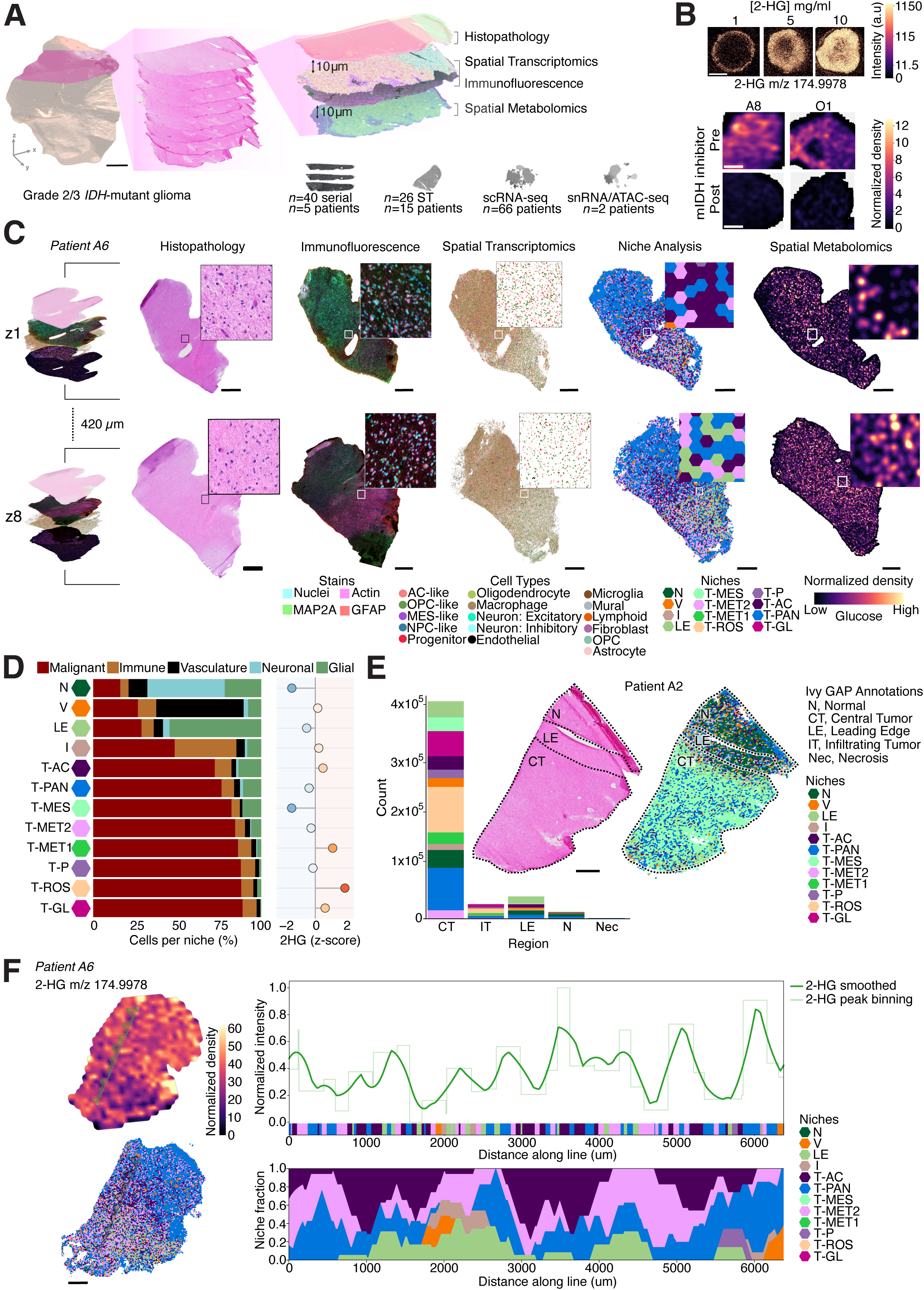
A multi-modal spatial atlas of *IDH*-mutant glioma. **A.** Representative overview of tissue sectioning strategy for deeply profiled patients, including micro-CT imaging. Eight multi-omic layers, represented by H&E stained sections, each contain serial sections of histopathology, ST with immunofluorescence and SM. Inset, sample and patient numbers for each modality of multi-omic atlas. Scale, 1 mm. **B.** 2-HG standard measured by SM. Above, intensity of 2-HG at m/z 174.9978 across increasing concentrations of 2-HG (1 mg/mL, 5 mg/mL, 10 mg/mL). Scale, 500 µm. Below, normalized density of 2-HG at m/z 174.998 in *IDH*-mutant glioma samples (A8 and O1) pre- and post-treatment with the mIDH inhibitor, safusidenib. Scale, 500 µm. **C.** Representative overview of spatial outputs of deeply profiled sample A6 at level 1 (z1) and level 8 (z8), 420 µm apart. Histopathology annotation, immunofluorescence, ST, GraphSAGE niches and SM of glucose are shown at each level. Scale, 1 mm. **D.** Proportion of major cell types present in each of the 12 GraphSAGE niches, with normalized abundance of 2-HG within each niche. **E.** Abundance of GraphSAGE niches in annotated Ivy GAP histopathology regions. Inset, representative section of Patient A2, with H&E (left) and niche distribution (right). Both sections overlaid with Ivy GAP pathology annotations, highlighting the LE region. Scale, 1mm. **F.** 2-HG metabolite density (above) and GraphSAGE niche distribution (below), with trajectory line of right inset overlaid in Patient A6. Scale, 1mm. From trajectory line, normalized 2-HG intensity across line distance. Bottom, GraphSAGE niche distribution across trajectory line.

### Tumor niches transcend anatomical organization

To identify recurring spatial niches that were conserved across patients, disease stages and treatment histories, we used the graph neural network algorithm GraphSAGE to perform transcript level analysis within 70 µm niches (**Figure 1C**). Across all ST samples, we identified twelve GraphSAGE spatial niches that maintained consistent spatial organization across serial sections (**Figure S1F, G**). These comprised four normal-cell (non-malignant) enriched and eight tumor-associated niches, designated according to their cell-type composition, transcriptional and metabolic features. Three niches corresponded to neural (N), vascular (V) and immune (I) microenvironments and were enriched for their respective normal cell populations (**Figure 1D**). Notably, each of these microenvironmental niches remained substantially infiltrated by malignant cells (>16% of cells), highlighting the known integration of tumor cells within normal brain tissue. Although V and I niches occupied only limited spatial areas (average contiguous size: V = 0.14 mm^2^, I = 0.03 mm^2^), both were identified in >75% of patient samples (**Figure S1H**), indicating that these microenvironmental ecosystems are conserved components of mIDH glioma architecture. We next evaluated the relationship of histopathological structures with the spatial niches (**Figure 1E, S1I, J**). Niches N and leading edge (LE) enriched with non-malignant cells generally mapped to their Ivy GAP anatomical features, however, most tumor-associated niches transcended traditional histological boundaries (**Figure 1E, S1I, J**).

Given that the production of 2-HG is the defining metabolic alteration of mIDH tumors, we next investigated how 2-HG was distributed across the tumor landscape. 2-HG spatial annotation was confirmed both by standards and its reduction in response to mIDH inhibition^17^ (**Figure 1B**). Amongst the niches associated with non-malignant cell composition, those anatomically aligned with non-tumor regions (N, LE) displayed significantly reduced 2-HG abundance, compared to the tumor disseminated V and I niches (**Figure 1D**). Variation of 2-HG abundance within the tumor-associated niches suggests intrinsic differences in the underlying spatial tumor metabolism (**Figure 1D, F, S1K**).

### Spatial niche identity is not determined by cellular composition alone

The eight tumor-associated niches were defined by distinct composition of malignant cell states, microenvironment and transcriptional activity (**Figure 2A, B, S2A**). To extend transcriptional characterization beyond the restricted spatial gene panel, we used CytoSPACE to map cells from our matched snRNA-seq data onto the ST data, providing transcriptome-wide expression profiles for each niche^25^ **(Table S3).** Pathway analyses compared transcriptional programs between niches while accounting for differences in cellular composition, thereby identifying pathways relatively enriched within each niche (**Figure 2B, S2B**). A pan tumor niche (T-PAN) represented the most prevalent tumor niche (average contiguous size: 6.2 mm^2^), occurring across all samples. This niche was characterized by a high cellular density (**Figure 2A**) and enrichment of canonical cancer-associated signaling pathways, including PDGF, RAS–MAPK, JAK–STAT, TGF-β and BMP signaling, relative to other niches (**Figure 2B**). We observed marked differences in compositional and transcriptional reprogramming between niches, even when in close proximity within samples. For example, the prevalence of an astrocyte-enriched tumor niche (T-AC, average contiguous size: 8.8 mm^2^) across samples was comparable to T-PAN (**Figure 2A**). However, this niche was characterized by substantially lower cell density and showed no enrichment for canonical cancer-associated signaling pathways (**Figure 2A**, **B**). Instead, T-AC was enriched for cholesterol biosynthesis and gap-junction signaling compared to other niches, features consistent with its enriched astrocytic composition (Chi-squared test against global astrocyte abundance, Pearson residual = 284.2, p-value < 0.001; **Figure 2B, S2A**). In addition to cellular and composition differences, we also observed elevated KEAP1/NRF2 pathway activity in T-PAN, T-ROS, named for its transcriptional oxidative stress response features, and T-GL, named for enrichment of the glia tumor compartment (CAMERA competitive gene set test^26,27^, log-fold change: T-PAN = 0.028, T-ROS = 0.048, T-GL = 0.062, false discovery rate (FDR): T-PAN = 0.027, T-ROS < 0.001, T-GL < 0.001). This highlights the spatial variation in oxidative stress response mechanisms. Thus, these data highlight complex spatial organization at the cellular and transcriptional levels.

**Figure 2.**
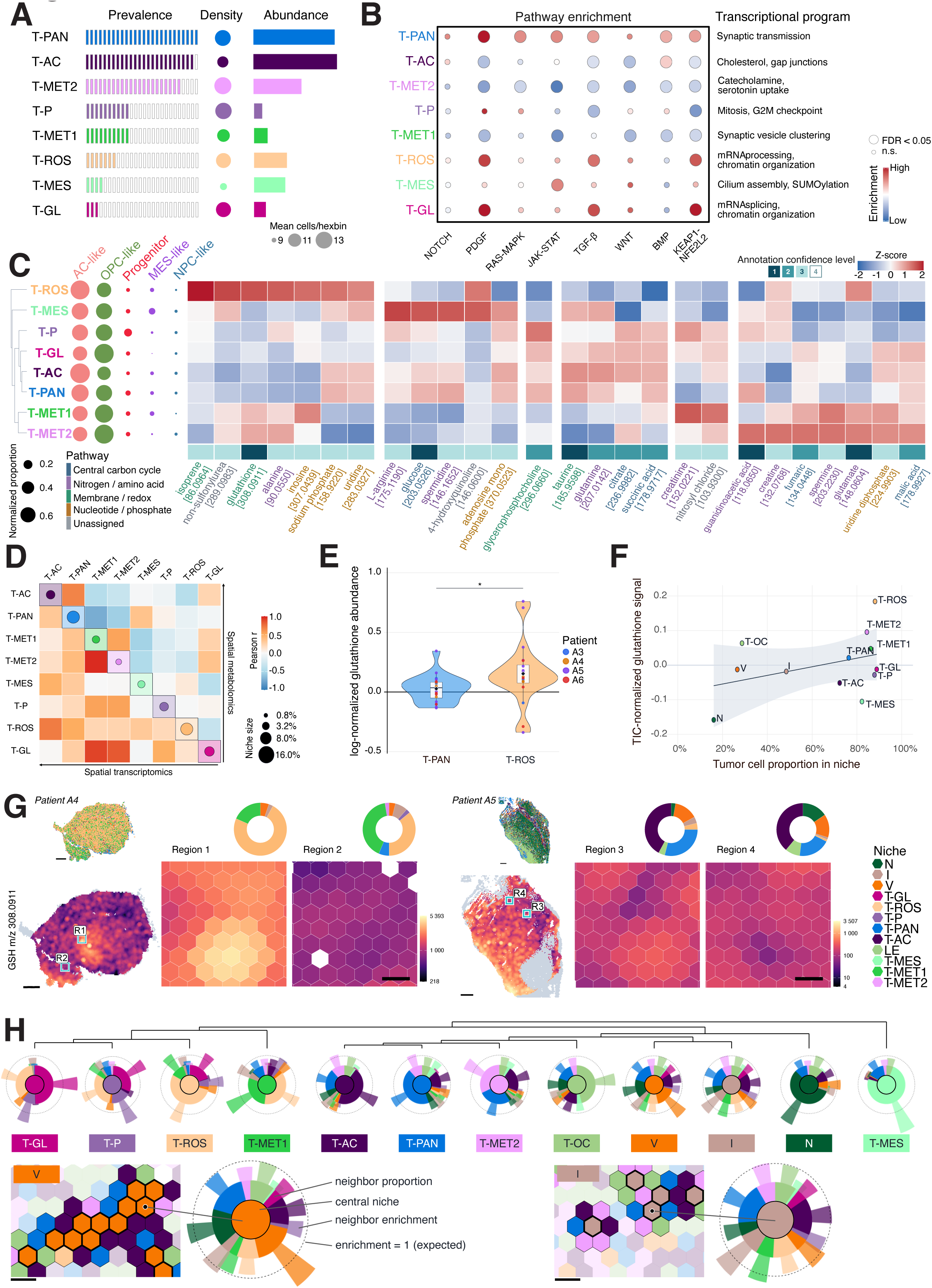
Tumor-associated niches are metabolically specialized. **A.** Spatial characteristics of tumor-associated niches identified from all ST samples. Prevalence, number of samples where niche occurs in at least 40 hexbins; density of cells within individual 70 µm hexbins for each niche; abundance of niche hexbins present across samples. **B.** Enriched pathways and semantic transcriptional programs as derived from CytoSPACE whole-transcriptome profile inference for each niche. **C.** Dendrogram of tumor-associated niches measured by normalized abundance of all metabolites identified in 75% of SM samples with abundance of annotated metabolites across tumor-associated niches. Represented with the normalized proportion of tumor cell-states per niche. **D.** Pearson r correlation of niches using their ST (lower triangle) and SM profiles (upper triangle). **E.** Log-normalized relative expression level of glutathione in T-PAN and T-ROS niches. **F.** Correlation of tumor cell proportion and normalized glutathione abundance in each niche. Shaded region represents 95 % confidence interval. **G.** Glutathione density across representative T-PAN- and T-ROS-enriched regions of patient A6 and A4, respectively. Scale, 1 mm. Left, sample niche distributions with zoomed-in region indicated. Scale, 100 µm. Centre, niche distributions of zoomed in regions. Right, glutathione density of zoomed in regions. **H.** Dendrogram of niche neighbors. Above, dendrogram of niche neighbors for each niche, calculated from similarity of niche neighbor proportions; with central circle indicating niche, middle ring displaying proportions of neighboring niches, and outer ring showing the square-root of the enrichment of niches among neighbors (segments over the dotted line represent enrichment). Below, representative regions of tissue for patient A6, with immediate neighbors of V and I niche hexbins respectively highlighted. Scale, 100 µm.

The observed niche heterogeneity prompted us to determine the link between cellular composition and metabolic microenvironments. To focus on reproducible metabolic features, we restricted our analysis to metabolite signals detected in all patients profiled with SM (as determined by the criteria of presence in > 75% of replicates from each patient) and clustered niches according to their resulting metabolic profiles (**Figure 2C, S2C, D, Table S2**). This revealed a metabolic organization that was distinct from differences in either malignant or non-malignant cellular composition (**Figure 2D**), indicating that niche metabolic identity was not simply determined by the cell populations present. Supporting the biological coherence of this organization, transcriptional and metabolic similarity between niches were correlated (**Figure S2E**). Two metabolically distinct niches (T-MET1 and T-MET2; average contiguous size: T-MET1 = 0.23 mm^2^; T-MET2 = 2.7 mm^2^; **Figure 2C**) showed no preferential enrichment of the canonical cancer-associated signaling pathways (**Figure 2B**). A tumor progenitor-enriched niche (T-P) was a densely cellular niche occupying a relatively small spatial footprint (2.9% of all niches; average contiguous size: 0.08 mm^2^; Chi-square test for progenitor enrichment, Pearson residual = 218.6, p-value < 0.001; **Figure 2A, S2A**). This niche also showed relative enrichment of pathways associated with cell-cycle progression, including regulation of mitotic nuclear division and the G2/M checkpoint (**Figure 2B, S2B**). A niche enriched for MES-like tumor cells (T-MES) was a rare niche, restricted to four individual samples (from 4 out of 15 patients), but expansive when present (average contiguous size: 25.7 mm^2^). Its enrichment of MES-like cells (Chi-square test, Pearson residual = 229.4, p-value < 0.001) was accompanied by a concordant relative decrease in 2-HG abundance (**Figure 2C, 1D, S2A**).

### Metabolic specialization reveals higher-order tumor organization

We next investigated level 1 annotated metabolites^28^ and their relative abundance across the tumor-associated niches (**Figure 2C, S1B**). T-PAN, T-AC, T-GL and T-P shared a core metabolic profile characterized by elevated tricarboxylic acid (TCA) cycle metabolites, including citrate and succinate, together with glutamine, a key carbon and nitrogen source. Taurine, the most abundant free amino acid in the brain and predominantly synthesized by astrocytes, was also elevated across these core tumor niches (**Figure 2C**). Significant depletion in taurine and divergence from the core tumor metabolism was observed in the T-MET1 and T-MET2 niches compared to T-AC (**Figure S2F**; Wilcoxon rank-sum test, p-values: T-AC versus T-MET2 = 0.0023; T-AC versus T-MET2 = 0.0642). T-MET1 and T-MET2 were further characterized by a dichotomy in guanidinoacetate (GAA) abundance. In T-MET2, GAA and creatine were abundant, while T-MET1 contained elevated creatinine, with lower GAA and creatine levels (**Figure 2C**), consistent with reduced 2-HG levels in T-MET2 compared to T-MET1 (**Figure 1D**), and reminiscent of impaired creatine synthesis and tumor-neuronal activity recently described in *IDH*-wildtype GBM^29^. The T-ROS niche exhibited a particularly distinctive metabolic signature. Reactive oxygen species (ROS) are produced in aerobic conditions and are increased under cellular stress^30^. Antioxidants are critical to neutralize ROS and maintain cell survival under deleterious conditions. The NRF2 pathway, significantly enriched transcriptionally in T-ROS (**Figure 2B**), is the master regulator of the antioxidant response, activated by oxidizing conditions and directly results in the production of glutathione. Glutathione was increased in abundance in T-ROS compared to the generalized T-PAN niche (**Figure 2E**; Wilcoxon signed-rank test, p-value = 0.035). This abundance was disproportionately increased relative to the average amount of glutathione given the tumor cell content (**Figure 2F**) and spatially distinct to the T-ROS niche (**Figure 2G**). These features suggest that T-ROS represents a metabolically distinct niche adapted to surviving cellular stress.

Analysis of niche adjacency revealed that spatial organization was not random but instead reflected higher-order organization (**Figure 2H**). The V and I niches showed no preferential association with specific tumor niches (mean absolute log-fold change of tumor niches: V = 0.83, I = 0.61, all other niches > 1.57), indicating that vascular support and immune interactions represent broadly shared compositional requirements. Consistent with this organization, hierarchical clustering based on niche adjacency closely mirrored clustering based on metabolite composition across tumor-associated niches (**Figure 2C, H**). T-MET1 and T-MET2 were the only pair to deviate from this trend, exhibiting strong metabolic similarity despite spatial segregation. Together, these findings suggest that higher-order tumor organization emerges through the interplay of local brain architecture, metabolic specialization and shared microenvironmental support.

### A *RIT2*+ intermediate malignant state is enriched in oligodendroglioma

To address tumor cell characterization, we combined our snRNA-seq data (n=32 samples from 22 patients with grade 2 and 3 mIDH glioma, **Table 1**) with publicly available grade 2 and 3 mIDH glioma datasets^6,7,17,31,32^ to generate a reference atlas spanning 101 samples from 66 tumors (45 astrocytomas, 21 oligodendrogliomas), comprising 574,870 nuclei and cells passing quality control (**Figure 3A, S3A, Table S4**). Nuclei and cells were integrated using scVI and annotated using marker gene expression (**Figure S3B**). In addition to the previously described canonical populations^33^, this analysis revealed a malignant state positioned between AC-like and OPC-like tumor programs (**Figure 3A, B**). These tumor cells were identified across all investigated datasets but one (**Figure S3C**), predominantly associated with oligodendroglioma (Wilcoxon rank-sum test, p-value < 0.001; **Figure 3C**), and had the same 1p/19q co-del status as the other malignant cells in oligodendroglioma (**Figure S3D**). Moreover, a distinct expression pattern was confirmed in publicly available bulk data (**Figure S3E, Methods**). Consistent with enrichment in oligodendroglioma, these cells were largely absent from held-out grade 4 mIDH astrocytomas of the same publicly available datasets (**Figure S3F**). This state was characterized by genes involved in neuronal signaling and cell-cell communication, such as highly specific expression of *RIT2* (**Figure 3D**), a marker of neuronal-like transcriptional programs^34^. The genes associated with *RIT2*+ tumor cells were not located in the 1p19q co-deleted regions that define oligodendrogliomas (**Figure 3D**). Corresponding to the genes associated with neuronal signaling, cells in the *RIT2*+ tumor cell state showed an enrichment of GABAergic signaling among all malignant populations (Wilcoxon signed-rank test, p-value < 0.001; **Figure S3G, H, Table S5**). Notably, several defining markers of these cells, including *RIT2* itself, followed developmental trajectories spanning both astrocytic and oligodendroglial lineages in the normal brain (**Figure S3I**). Despite these features, *RIT2*+ tumor cells remained transcriptionally distinct from both a recently described OPC-GABA population^35^ (**Figure S3J**) and an undifferentiated malignant population^6^ (**Figure S3K**).

**Figure 3.**
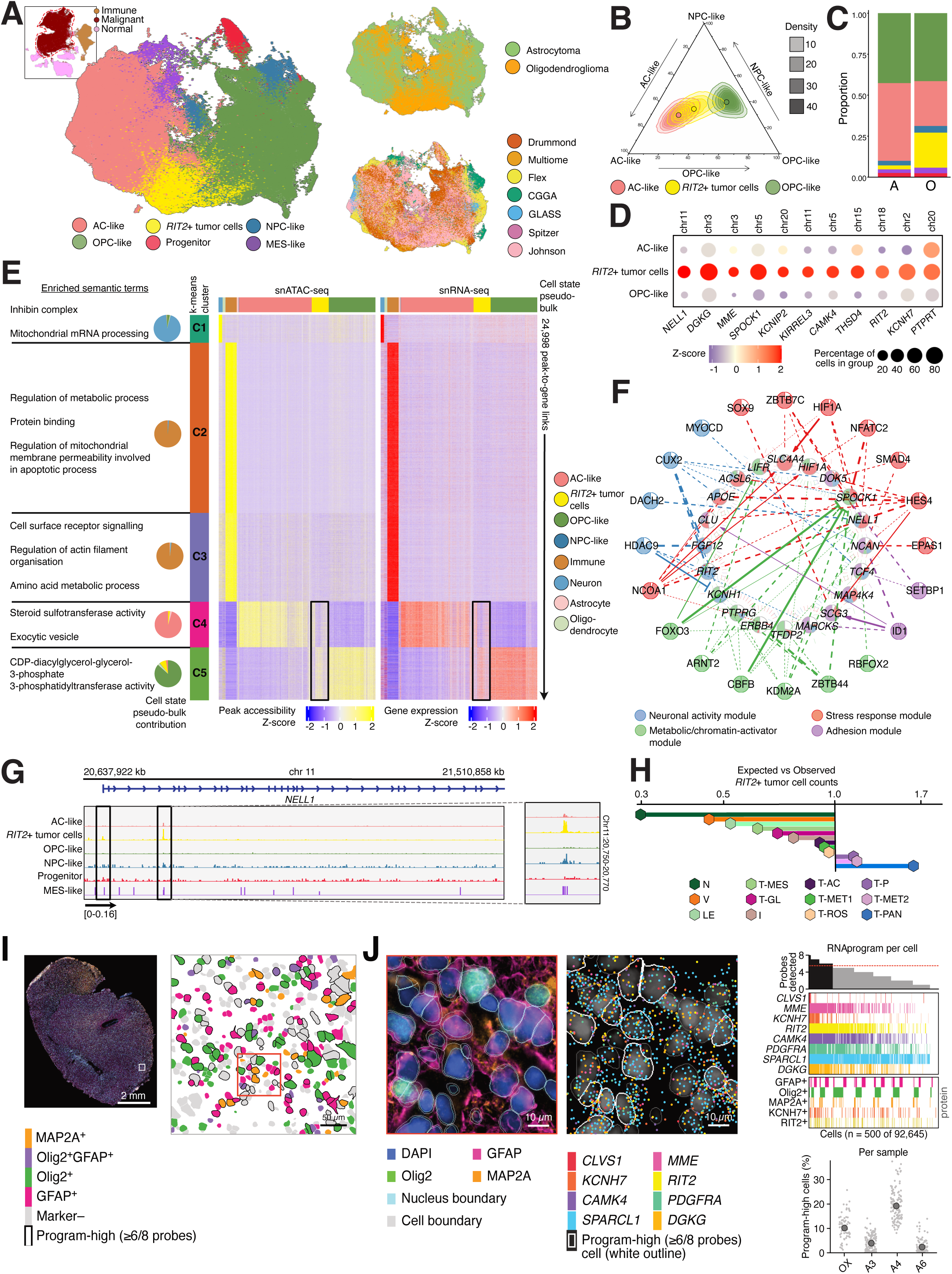
A *RIT2*+ intermediate malignant state is enriched in oligodendroglioma. **A.** UMAP depicting reference atlas of 574,870 cells and nuclei spanning 101 samples from 66 patients with grade 2 or 3 mIDH glioma. Left: annotated UMAP clusters of tumor cell states only accompanied by inset depicting all malignant and non-malignant cells; top right: glioma subtype; bottom left: dataset source. **B.** Ternary plot depicting transcriptional glioma programs; AC-like, OPC-like and NPC-like, mapped with AC-like, *RIT2*+ and OPC-like tumor cells, outlined circles represent the mean position of each cell type. **C.** Proportion of each malignant cell state (key in A) in astrocytoma (A; n = 45) and oligodendroglioma (O; n = 21) tumors. **D.** Dot plot of z-score of log-normalized expression of candidate *RIT2*+ tumor cell markers in AC-like, OPC-like and *RIT2*+ tumor cells. Chromosome location of each marker indicated above. **E.** Heatmap of chromatin accessibility (left) and gene expression (right) of 24,998 putative cis-regulatory peak-to-gene associations (rows) across cell state pseudo-bulks of snRNA-seq and snATAC-seq nuclei, clustered into k-means clusters. Enriched semantic terms and cell state composition shown in pie charts for clusters. **F.** Annotated NeighbourNet^36^ meta-network 3 enriched in *RIT2*+ tumor cells of oligodendroglioma samples. Inner layer: target genes; outer layer: central transcription factors. Edges and their thickness indicate evidence of interaction: none (no edge), significant co-expression only (dashed), significant co-expression supported by prior evidence (solid, with arrowhead for activation or barhead for repression). Targets and transcription factors colored by proportion module occupancy. **G.** Chromatin accessibility track of *NELL1* locus by tumor cell state. Right inset: zoomed region of differentially accessible peak in *RIT2*+ tumor cells. **H.** Enrichment of *RIT2*+ tumor cell localization across GraphSAGE niches. Ratio of observed to expected cell counts of *RIT2*+ cells within each niche, based on overall niche abundance. **I.** COMET sequential IF and RNAscope of patient A4. Left: Section overview (DAPI, GFAP, Olig2, MAP2A; scale bar, 2 mm). Right: Segmented cells in the boxed 350 µm field colored by IF phenotype; cells detecting >= 6 of 8 program probes (program-high) outlined. Scale bar, 50 µm. **J.** Enhanced area outlined in I. Right: IF with nucleus and cell segmentation. Middle: RNAscope transcripts of the 8 program probes with program-high cells outlined. Scale bars, 10 µm. Right: Probes detected per cell for 500 random cells, with protein detection (GFAP, Olig2, MAP2A, KCNH7, RIT2) for the same cells, and % program-high cells per sample per 0.57 mm block.

To investigate mechanisms that regulate gene expression in the *RIT2*+ tumor cells, we analyzed matched chromatin accessibility and gene expression profiles generated using simultaneous single nuclei RNA and chromatin accessibility (snATAC) sequencing from three grade 2 mIDH gliomas (**Figure S4A, B, Table S6**). Consistent with their intermediate transcriptional profile, *RIT2*+ tumor cells occupied an analogous location between AC-like and OPC-like populations in chromatin accessibility space (**Figure S4C**). With these data we identified open chromatin regions associated with genes across all nuclei. Putative cis-regulatory elements were clustered by their activity patterns across both chromatin and transcriptome into regulatory modules (**Figure 3E**). Regulatory modules enriched in AC-like cells were associated with secretory pathways, whereas OPC-like modules were enriched for phospholipid biosynthesis pathways. Notably, *RIT2*+ tumor cells were the only malignant population to exhibit accessible chromatin associated with both AC-like and OPC-like regulatory programs, whereas canonical AC-like and OPC-like populations largely showed accessibility restricted to their respective lineage-associated modules^33^. These findings suggest that *RIT2*+ tumor cells retain regulatory features of both tumor programs rather than representing simple extensions of either cell state.

We therefore sought to determine whether *RIT2*+ tumor cells were additionally characterized by a distinct regulatory architecture that could explain their unique transcriptional identity. Applying NeighbourNet^36^, we identified a unique program representing *RIT2*+ tumor cells across the oligodendroglioma tumors (**Figure S4D, S4E**). The identity of this distinct regulatory network was driven by neuronal, metabolic, stress and adhesion modules (**Figure 3F, Table S7**). Within the stress response module, the *RIT2*+ tumor cell marker *NELL1* was highly connected. Notably, *NELL1* is more highly expressed in OPCs than in mature oligodendroglial and astrocytic lineages in the normal developing brain, consistent with its role in differentiation (**Figure S4F**). Importantly, some predicted regulatory interactions were independently supported by snATAC-seq data. For example, motif analysis of accessible chromatin regions identified the Notch-responsive HES-repressors bound at the N-box (CACNAG; 63.3% N-box positive) in the *NELL1* promoter. This indicates competency for HES4 binding, and together with concomitant *NELL1* and *HES4* expression in *RIT2*+ tumor cells, implicates Notch signaling in the regulation of *NELL1* and differentiation within the stress-adaptive program (**Figure 3G**).

We next investigated the spatial organization of *RIT2*+ tumor cells. Prediction of *RIT2*+ tumor cell identity within the ST showed their enrichment within the T-PAN niche and frequently localized adjacent to both AC-like and OPC-like tumor populations (**Figure 3H, S4G, H, Methods**). However, despite their distinct molecular features, the local cellular environments surrounding *RIT2*+ tumor cells were highly similar to those observed around AC-like and OPC-like populations, with no evidence for a unique immediate neighborhood composition (**Figure S4H**). To independently confirm the presence of this population in situ, we combined multiplexed IF with RNAscope targeting eight genes associated with the intermediate transcriptional program (**Figure 3I, J, S4I**). Program-high cells were identified across four independent tumors, providing orthogonal validation of this transcriptional identity. Together, these findings identify a spatially interspersed *RIT2*+ tumor cell population, revealing greater phenotypic heterogeneity in mIDH glioma, particularly oligodendroglioma, than previously appreciated.

### Niche specialization of the myeloid compartment

Our analysis demonstrated myeloid cells were detectable across all niches including specific enrichment within the I niche (**Figure S2A**). Given the central role of myeloid cells in the immunosuppressive glioma microenvironment, and their interplay to metabolic cues such as 2-HG, we characterized their spatial identity. In our snRNA-seq data, the myeloid population was composed of resident microglia as well as infiltrating monocytes, macrophages and dendritic cells (**Figure 4A, S5A**). As these broad cell annotations do not fully capture functional immune states, we next classified myeloid cells using a recently described immunomodulatory framework^37^. This approach allows myeloid cells to be grouped into four functional programs that define inflammatory and immunosuppressive states independent of conventional myeloid cell identity (**Figure 4B**). Grade 2 oligodendrogliomas and astrocytomas both occupied predominantly inflammatory regions of the immunomodulatory landscape, while grade 3 astrocytomas and oligodendrogliomas shifted toward the center of the program space. GBM were more immunosuppressive, consistent with progressive acquisition of immunosuppressive features during malignant progression^37^ (**Figure 4C, S5B, C**). The four immunomodulatory programs were originally defined using whole-transcriptome single-cell data, however targeted ST panels capture only a subset of the genes comprising these signatures (**Table S8**). As a result, direct application of the original program scores leads to systematic distortion of immunomodulatory state estimates preventing reliable spatial mapping (**Figure 4D, S5D**). To overcome this limitation, we developed a non-linear calibration approach that learns how targeted gene panels distort immunomodulatory program scores by simulating panel-restricted measurements from whole-transcriptome snRNA-seq data (**Figure 4D, Methods**). Applying this learned correction to the panel-restricted measurements accurately reconstructed the underlying whole-transcriptome program distributions, restoring correlations between gene expression and program localization, and significantly reducing the distance between ground truth and observed placements (median distance, restricted: 18.2 units; adjusted: 10.0 units; paired t-test, mean decrease of 8.97 units, p-value < 0.001) (**Figure 4D, S5D-G**). We validated the applicability of this approach in three independent GBM samples, in which the methodology successfully recovered the underlying immunomodulatory program distributions (**Figure S5H-K, Table S9**).

**Figure 4.**
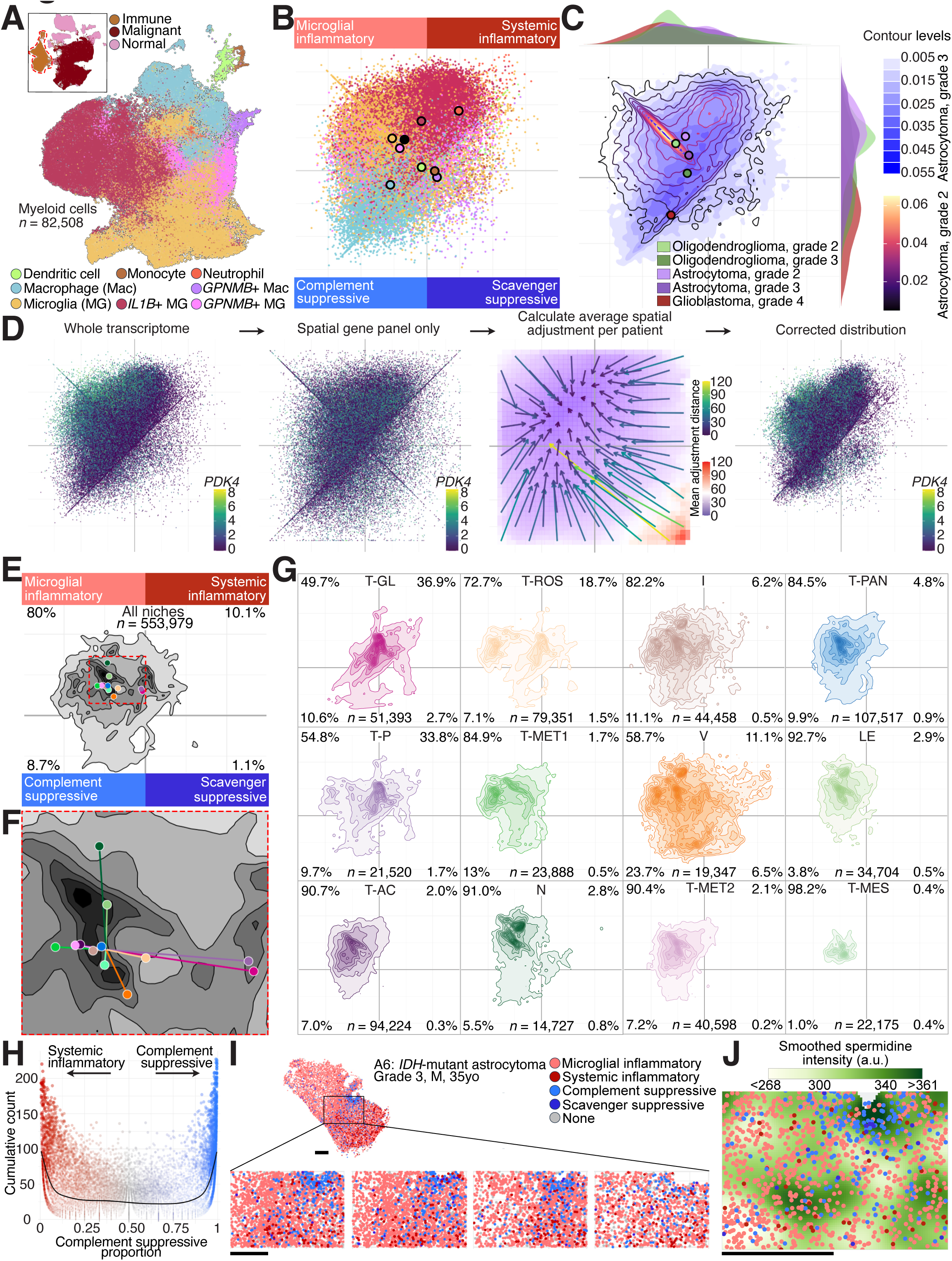
Myeloid immunomodulatory programs partition tumors into inflammatory and immunosuppressive territories with altered spermidine abundance. **A.** Myeloid annotation of snRNA-seq cohort. UMAP of snRNA-seq myeloid cells, colored by cell type annotation. Inset, UMAP of snRNA-seq cohort, with immune clusters outlined. **B.** Immunomodulatory program enrichment of snRNA-seq myeloid cells, colored by cell type annotation. Outlined circles represent the mean position of each cell type, with the solid black circle representing the global mean position. **C.** 2D density of immunomodulatory program distribution of snRNA-seq astrocytoma samples, split into grade 3 (blue gradient) and grade 2 (‘magma’ gradient) samples. Outlined circles represent the mean position of each classification, including grade-stratified oligodendroglioma and additional glioblastoma samples. 1D density of classifications shown along both axes. **D.** Immunomodulatory program calibration strategy. From left to right: log-normalized expression of *PDK4* in snRNA-seq myeloid cells distributed by immunomodulatory program score using whole-transcriptome profiles; *PDK4* expression distributed by immunomodulatory program score using ST restricted gene profiles; Average spatial adjustment to correct distribution of ST restricted program score to whole-transcriptome program score distributions; *PDK4* expression distributed by immunomodulatory program score using ST restricted gene profiles after calibration (**Methods**). **E.** 2D density of ST myeloid cell immunomodulatory program distribution, after calibration using strategy in (D). Proportion of cells in each quadrant displayed, and mean positions of niches represented by outlined circles. **F.** Zoomed-in region of (E) depicting mean displacement of niche myeloid programs from global mean location. **G.** Niche-specific 2D densities of ST myeloid cell immunomodulatory program distribution, with proportions of cells in each quadrant displayed. **H.** Comparison of overall number of ST myeloid cells within systemic inflammatory and complement suppressive programs within a region, and the proportion of complement suppressive cells within these. Only ST myeloid cells with systemic inflammatory and complement suppressive programs scores of ≥ 20 were included in analysis. Measured regions were 1 mm by 1 mm across all ST samples. **I.** Representative region of sample A6, with myeloid cells colored by immunomodulatory program (programs scores of ≥ 20; or for scavenger immunosuppressive, ≥ 5). Region shown across 4 serial ST layers. Scale, 1mm. **J.** Smoothed abundance density of spermidine in region of sample A6 shown in (I), overlaid with myeloid cells colored by immunodmodulatory program shown in (I). Scale, 1mm.

Using our calibration approach, we inferred immunomodulatory program usage for individual cells in the ST dataset, enabling spatial reconstruction of functional myeloid states despite the limited gene panel (**Figure 4E-F**). Consistent with their shared roles in immune surveillance and tissue homeostasis, the I and V niches contained the greatest diversity of immunomodulatory programs, with the largest immunosuppressive program proportions residing in the V niche (**Figure 4E-G**). In contrast, immunosuppressive states were infrequent in the tumor-associated niches, but enriched in the spatially associated T-ROS, T-GL and T-P niches (**Figure 4G**). Notably, tumor niches that preferentially neighbored one another also shared similar immunomodulatory program compositions, indicating that immune states are coordinated across higher-order tumor ecosystems rather than independently specified within individual niches (**Figure 4F, G, 2H**).

Investigating the spatial relationship between systemic inflammatory and complement suppressive programs across the broader cohort, we found that they occupied spatially distinct territories and rarely co-localized within the same local neighborhoods (**Figure 4H, I**). This behavior was unlike what was observed for other program pairs (**Figure S5L**). Rather than representing independent axes of myeloid activation, systemic inflammatory and complement suppressive programs defined mutually exclusive immune microenvironments that partitioned the tumor into inflammatory and immunosuppressive regions (**Figure S5M**). These distinct immune territories were also associated with differential spermidine enrichment (t-test, log-fold change=−0.528, inflammatory versus immunosuppressive; **Figure 4J**). This observation of spermidine enrichment in immunosuppressive regions is consistent with the established role of spermidine as a supporting factor in immunosuppressive myeloid and lymphoid function^38^. This compartmentalized organization suggests that these programs represent alternative spatial immune ecosystems.

### Expansion of a metabolically specialized niche following treatment

Given the limited understanding of how the tumor microenvironment evolves in mIDH glioma, we investigated its response to standard-of-care chemoradiation and emerging mIDH inhibitor therapies. Previous studies investigating the impact of treatment in mIDH glioma in bulk or cell specific contexts failed to identify an intrinsic response to chemoradiation or mIDH inhibition at relapse^32,39^. Longitudinal data pairs were profiled in the snRNA-seq cohort, consisting of six samples paired with no adjuvant treatment (surgery 1 and surgery 2), twelve samples paired with mIDH inhibitor treatment from a perioperative clinical trial of safusidenib^17^, and twelve samples paired with adjuvant alkylating chemotherapy and/or radiotherapy (chemoradiotherapy)^6^.

Given initial insights into inflammation at recurrence^6^, we first investigated the myeloid compartment phenotypes. Using longitudinal pairs with no adjuvant treatment as a baseline, we found the impact of surgery was negligible on the phenotype of myeloid cells (**Figure 5A**). Similarly, the response to mIDH inhibition, though reported to increase macrophage numbers in the tumor^17^, resulted in similar microglial inflammatory phenotypes as the untreated counterparts. In contrast, chemoradiation resulted in a shift to a suppressive phenotype, similar to the effect of increased histological grade (t-test, mean 13.5 unit decrease along y-axis post-chemoradiation, p-value < 0.001; **Figure 5A, 4C**). This significant shift, predominantly to the scavenger suppressive phenotype (t-test, mean 13.9 unit shift towards scavenger suppressive quadrant post-chemoradiation, p-value < 0.001) represents a forced evolution of the myeloid compartment directed by treatment, irrespective of the tumor grade.

**Figure 5.**
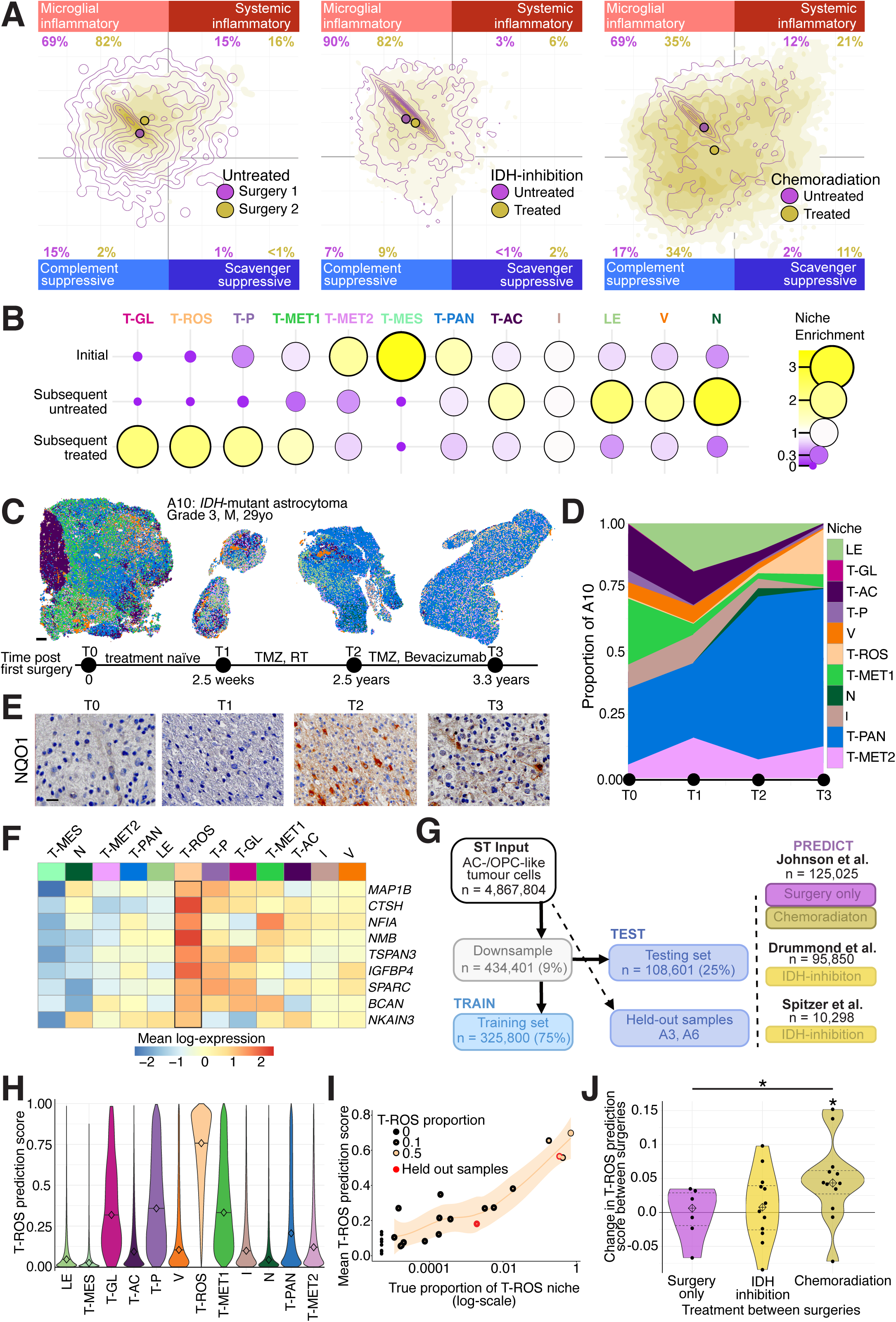
A glutathione-rich antioxidant niche expands after chemoradiotherapy. **A.** Density of snRNA-seq myeloid cells distributed by immunomodulatory program score, stratified by treatment and surgical timepoint. Purple contours display first surgical timepoint; gold gradient displays second surgical timepoint. From left to right: samples without treatment between surgeries (surgery only; n = 12); samples with mIDH-inhibition treatment between surgeries (n = 24); samples with chemotherapy and/or radiotherapy treatment between surgeries (n = 24). **B.** Enrichment of ST niches within samples from initial and subsequent surgeries, with subsequent surgeries further stratified by prior treatment. **C.** GraphSAGE niche distribution of longitudinal samples (T0 to T3) from patient A10. Intervening treatment listed on the timeline. Scale, 1 mm. **D.** Niche proportion transitions between longitudinal samples of patient A10. **E.** Representative immunohistochemical staining of NQO1 abundance in longitudinal samples of patient A10. Scale, 20 µm. **F.** Mean log-expression of T-ROS-associated genes across all niches. **G.** Schematic for logistic regression classifier for prediction of T-ROS-associated tumor cells in snRNA-seq data, using downsampled ST data for training. **H.** Violin plot of T-ROS prediction score in ST testing set (n = 108,601) per niche, with median indicated. **I.** Relationship between proportion of T-ROS niche within ST samples, and overall mean T-ROS prediction score in that sample. Held out samples are highlighted in red. **J.** Violin plot of change in mean T-ROS prediction between surgical timepoints, stratified by treatment between surgeries: surgery only, mIDH inhibition, or chemotherapy and/or radiotherapy. Asterisks indicate p-values < 0.05 for Wilcoxon rank-sum exact tests, either for individual stratification (null hypothesis: mean change is not significantly different to zero) or between stratifications (null hypothesis: difference between mean changes is not significantly different to zero).

To determine the impact of treatment on the tumor ecosystems, we calculated the frequency each niche was found in the longitudinal samples, from first surgery to second with or without adjuvant treatment (**Figure 5B**). Strikingly, T-ROS, T-P and T-GL niches were enriched in the treated microenvironments (individual Chi-squared tests against all untreated conditions combined, all p-values < 0.001). T-ROS, the most abundant niche of the three (**Figure 2A**), was found to have disproportionately high glutathione (**Figure 2E**) and elevated 2-HG compared to other niches (**Figure 1D**). This marked spatial accumulation of glutathione in the T-ROS niche (**Figure 2G**), supports its designation as a redox-associated ecosystem adapted to buffer oxidative stress, consistent with its emergence following treatment (**Figure 5B**). This pattern was exemplified by one longitudinally profiled patient with grade 3 astrocytoma, in whom the T-ROS niche was absent at diagnosis, remained rare following surgery alone, and expanded substantially after chemoradiotherapy and subsequent bevacizumab treatment (**Figure 5C, D**). Furthermore, consistent with the enhanced KEAP1/NRF2 pathway enrichment in the T-ROS niche (CAMERA competitive gene set test, log-fold change = 0.048, FDR < 0.001), this patient displayed accumulation of the NRF2 biomarker NQO1 as the T-ROS niche expands (**Figure 5E**). Together, these findings suggest that therapeutic pressure is associated with expansion of a metabolically specialized niche characterized by glutathione accumulation and enhanced antioxidant capacity.

To determine whether the T-ROS niche could be identified in the longitudinal snRNA-seq datasets, we first asked whether AC/OPC-like tumor cells residing within T-ROS exhibited a distinct transcriptional program. Despite sharing the same broad malignant cell states with other tumor niches, T-ROS-associated cells showed a reproducible set of differentially expressed genes, indicating that residence within this niche is accompanied by a detectable transcriptional signature (**Figure 5F**). We therefore trained a logistic regression classifier to distinguish AC/OPC-like cells within T-ROS from those in other tumor niches, using spatially annotated cells for training and held-out cells and samples for validation (**Figure 5G**). The classifier accurately identified T-ROS-associated cells in held-out spatial data (median prediction score: 0.69; 71% of T-ROS cells with prediction score ≥ 0.5) (**Figure 5H, S6A-F**). Elevated prediction scores were also observed in T-P, T-MET1 and T-GL (median prediction scores: 0.5, 0.32, and 0.56, respectively), suggesting partial sharing of the T-ROS transcriptional program with other treatment-associated tumor niches. Importantly, the mean predicted T-ROS signature score in ST samples closely correlated with their measured T-ROS niche abundance (**Figure 5I**). On a more granular level, hexbins with stronger underlying T-ROS signatures had increased prediction scores (correlation = 0.80, p-value < 0.001), which drove the elevated prediction scores of closely associated regions of niches T-P, T-MET1 and T-GL (**Figure S6G-H).**

We next applied the classifier to three longitudinal single-cell datasets to determine whether cells with the T-ROS signature emerge preferentially following specific therapeutic exposures. Overall, cells carrying the T-ROS signature represented only a small fraction of the AC/OPC-like population (median proportion of cells with a prediction score ≥ 0.5 = 7%). The mean T-ROS signature prediction score remained unchanged in longitudinal tumors undergoing surgery alone or following mIDH inhibition treatment (Wilcoxon rank-sum exact test, surgery alone: p-value = 1.0; mIDH inhibition treatment: p-value = 0.57; **Figure 5J**). In contrast, the T-ROS signature was significantly increased following chemoradiation treatment (Wilcoxon rank-sum exact test, p-value = 0.034; **Figure 5J**). Together with the spatial enrichment of T-ROS in recurrent treated tumors in our spatial cohort, these findings indicate that the T-ROS program is specifically associated with exposure to chemoradiation rather than recurrence or treatment more generally.

## DISCUSSION

By integrating spatial metabolomics, transcriptomics and histology across aligned tissue sections of grade 2 and 3 mIDH glioma tumors, we identified metabolic specialization that could not be explained by cellular composition or anatomical regions alone. Previous studies have emphasized the contributions of normal brain anatomy, vascularization, and hypoxia to glioma architecture^9,40^, while spatial multi-omic investigations in GBM have demonstrated how local metabolic and inflammatory cues shape malignant cell states and their organization^41^. Extending these observations to grade 2 and 3 mIDH gliomas, we identified twelve conserved spatial niches with distinct metabolic identities, whose metabolic similarity was reflected in their spatial adjacency. Together, these findings suggest that spatial organization in grade 2 and 3 mIDH gliomas, where extensive hypoxia and necrosis are less prominent, is shaped by the interplay of cellular identity, metabolic zonation, and local tissue interactions rather than a single dominant organizing principle.

Our identification of a distinct *RIT2*+ malignant population preferentially associated with oligodendrogliomas reveals a difference between the two mIDH glioma subtypes historically considered to share broadly similar malignant cell-state compositions^6,8^. Largely absent from WHO grade 4 disease, this population challenges prevailing notions of aggressive cellular features. It occupied an intermediate transcriptional and epigenetic state between AC-like and OPC-like cells and exhibited expression of genes associated with regulatory plasticity and neuronal signaling, features typically associated with cellular adaptability and proliferation in glioma^6,42–44^. Whether this population represents a reservoir of cellular plasticity or a differentiation-restricted state with limited malignant potential remains unclear. Its preferential association with lower-grade disease raises the possibility that these features do not necessarily confer greater aggressiveness but may instead, at times, reflect a constrained developmental state.

Longitudinal studies have established that mIDH glioma progression involves expansion of myeloid cell infiltration^45,46^, progressive loss of DNA methylation^15^ and treatment-associated hypermutation^47^. These changes occur against a background of broadly conserved malignant cell identities^6^. Our findings indicate that additional treatment-specific adaptations at progression become apparent when tumors are examined through the lens of metabolic zonation. We identified a rare niche characterized by glutathione accumulation and NRF2-associated antioxidant programs, which expanded following treatment. Although its limited extent may explain why this program has been overlooked in previous analyses, its transcriptional signature enabled its detection in independent longitudinal single-cell cohorts, where it increased following chemoradiation but not mIDH inhibition or surgery alone. Consistent with previous *in vitro* studies implicating NRF2 activation and glutathione-dependent antioxidant defenses in resistance of alkylating chemotherapy^48,49^, our findings identify redox adaptation as a spatially restricted component of treatment-associated evolution. The broader relevance of this pathway is underscored by studies in non-small-cell lung cancer, where activating mutations in the NRF2 pathway promote metabolic reprogramming, tumorigenesis, and remodeling of the immune microenvironment^50,51^. However, because samples were collected after rather than during treatment, whether the presence of this niche following treatment reflects the expansion or resolution of a treatment-induced metabolic adaptation, and to what extent these processes reverse following treatment withdrawal, remains unclear.

Together, these findings establish spatial metabolic specialization as an important dimension of glioma heterogeneity and adaptation. More broadly, our integrated atlas provides a framework for investigating tissue organization across spatial scales and molecular modalities in mIDH glioma. By profiling and aligning serial tissue sections across multiple depths, our approach lays the foundation for three-dimensional multi-omic reconstruction of tumor architecture, while computational methods that recover functional programs from restricted transcriptomic panels enable deeper characterization of spatially resolved cellular states. Together, these advances provide a resource for investigating metabolic and microenvironmental interactions that would otherwise remain difficult to resolve.

## ACKNOWLEDGMENTS

We thank S. Stylli for expert curation of the Royal Melbourne Hospital Neurosurgery Brain and Spine Tumour Tissue Bank, R. Linklater and R. Huynh for expert study management, M. Bisignano for assistance at the RMH Anatomical Pathology Department, E. Tsui for assistance at the WEHI Advanced Histotechnology Facility, C. Anttila, L. Ling and R. Mishi for assistance at the WEHI Advanced Genomics Facility and P. Lam, E. Watson and L. Xing for assistance at the Centre for Dynamic Imaging. We thank L. Smyth and E.D. Hawkins for critical review. This study was made possible and financially supported in part through the authors’ membership of the Brain Cancer Centre, through support from Carrie’s Beanies 4 Brain Cancer, through the Victorian State Government Operational Infrastructure Support and Australian Government NHMRC Independent Research Institutes Infrastructure Support Scheme (IRIISS) and the Victorian Government under the Brain Perioperative Clinical Trial Program (Brain-POP). Further support included funding from the Venture Grants Scheme administered by Cancer Council Victoria (VG2022 to S.A.B., S.F., J.R.W. and M.J.M.), the NHMRC (GNT2033815 to J.R.W.), the Medical Research Future Fund (MRF2019448 to K.J.D, S.A.B., S.F. and J.R.W), Victorian Cancer Agency Mid-Career Research Fellowship (MCRF22003 to S.A.B.), Cancer Council Victoria Mid-Career Research Fellowship (2025/CRFV0072 to S.F.), WEHI Page Betheras Award (S.A.B.), WEHI IPSI Scholarship (O.E.F.), WEHI Johnson PhD Scholarship (J.J.D.M.), Australian Government Research Training Program Scholarship (J.J.D.M., T.L., and M.S.), Dine for a Cure Legacy Scholarship (M.S.), CSL Translational Data Science Scholarship (J.J.D.M.), Bartlett Foundation support (J.K.), Mark Hughes Foundation Fellowship (S.P), Gregg Symons Scholarship for Brain Cancer Research (O.E.F.) and Perpetual IMPACT, William Angliss and Norman Ann and Graeme Atkins philanthropic programs. The funders had no role in study design, data collection and analysis, decision to publish or preparation of the paper.

## AUTHOR CONTRIBUTIONS

Conceptualization, S.A.B., S.F., and J.R.W.; Data curation, J.J.D.M., J.K., T.L., M.S., O.E.F., B.K., L.R., and S.J.R.; Formal analysis, J.J.D.M., J.K., T.L., M.S., O.E.F., B.K., L.R., M.L., and S.J.R.; Funding acquisition, S.A.B., S.F., J.R.W., K.J.D.; Investigation, J.K., T.L., A.L.V., J.F.P., Z.M., S.P., K.J.D., J.J., S.A.C., V.K.N., and B.N.; Methodology, S.F., J.J.D.M., J.K., T.L., and M.S.; Project administration, S.A.B., S.F., J.K., and J.R.W.; Resources, S.A.B., S.F., and J.R.W.; Software, J.J.D.M., J.K., T.L., and M.S.; Supervision, S.A.B., S.F., J.R.W., M.L., D.M.A., and M.J.M.; Validation, J.K.; Visualization, J.J.D.M., J.K., T.L., and M.S.; Writing – original draft, S.A.B., S.F., J.J.D.M., J.K., T.L., and M.S.; Writing – review & editing, all authors.

## DECLARATION OF GENERATIVE AI AND AI-ASSISTED TECHNOLOGIES IN THE WRITING PROCESS

During the preparation of this work, the authors used Claude (Anthropic) and ChatGPT (OpenAI) to edit and restructure the manuscript into Cell format. After using this tool, the authors reviewed and edited the content as needed and take full responsibility for the content of the publication.

## SUPPLEMENTAL INFORMATION

Document S1. Figures S1–S6 and Tables S1–S10

## STAR METHODS

### KEY RESOURCES TABLE

| REAGENT or RESOURCE | SOURCE | IDENTIFIER |
|---|---|---|
| **Antibodies** |  |  |
| Rabbit monoclonal anti-GFAP (clone E4L7M) | Cell Signaling Technologies | Cat# 80788; RRID: AB_2799963 |
| Chicken polyclonal anti-MAP2A | Thermo Fisher Scientific | [Cat# PA1-10005; RRID: AB_1076848] |
| Rabbit monoclonal anti-MAP2 (clone EPR19691) | Abcam | Cat# ab183830; RRID: |
| Rabbit monoclonal anti-SOX9 (clone 7H13L8) | Thermo Fisher Scientific | Cat# 702016; RRID: |
| Mouse monoclonal anti-GFAP (clone GA5) | Cell Signaling Technology | Cat# 3670; RRID: |
| Mouse monoclonal anti-IDH1 R132H (clone H09) | Dianova | Cat# DIA-H09; RRID: |
| Rabbit monoclonal anti-OLIG2 (clone E6G6Q) | Cell Signaling Technology | Cat# 65915S; RRID: |
| Rabbit polyclonal anti-KCNH7 | Alomone Labs | Cat# APC-112-200; RRID: |
| Rabbit polyclonal anti-CLVS1 | Thermo Fisher Scientific | Cat# OSC00321W-100UL |
| Rabbit polyclonal anti-NQO1 | Abcam | Cat# ab34173; RRID: |
| Rabbit polyclonal anti-RIT2 | Thermo Fisher Scientific | Cat# PA5-144175; RRID: |
| Goat anti-rabbit IgG (H+L), Alexa Fluor 594 | Thermo Fisher Scientific | Cat# A11039; RRID: AB_2534096 |
| Goat anti-chicken IgY (H+L), Alexa Fluor 488 | Thermo Fisher Scientific | [Cat# A48265; RRID: AB_2895299] |
| **Biological samples** |  |  |
| Human IDH-mutant glioma tissue, WHO CNS grade 2-3 (deeply profiled and extended spatial cohorts) | Royal Melbourne Hospital Neurosurgery Brain and Spine Tumour Tissue Bank | Table 1; Melbourne Health Ethics # 2020.214 |
| Paired pre- and post-safusidenib IDH-mutant glioma samples | Drummond et al.<sup>7</sup> | ClinicalTrials.gov: NCT05577416 |
| **Chemicals, peptides, and recombinant proteins** |  |  |
| Alexa Fluor 647 phalloidin | Thermo Fisher Scientific | Cat# A22287; RRID |
| DAPI | Thermo Fisher Scientific | Cat# D1306; RRID: AB_2629482 |
| Paraformaldehyde 16% aqueous solution, EM grade | Electron Microscopy Sciences | Cat# 15710 |
| Methanol, for HPLC | Millipore Sigma | Cat# 34860 |
| Ethyl alcohol, pure (200 proof, anhydrous) | Millipore Sigma | Cat# E7023-500ML |
| PBS (10X), pH 7.4, RNase-free | Thermo Fisher Scientific | Cat# AM9624 |
| 10% Tween 20 | Bio-Rad | Cat# 1662404 |
| TE buffer, Tris-EDTA, 1X, pH 8.0 (nuclease-free) | Fisher Scientific | Cat# BP24731 |
| Nuclease-free water | Thermo Fisher Scientific | Cat# AM9937 |
| Normal goat serum | Abcam | Cat# ab138478 |
| Fluoromount mounting medium | Proscitech | Cat# EMS17984-25 |
| α-Cyano-4-hydroxycinnamic acid (CHCA) MALDI matrix | Sigma-Aldrich | Cat# C2020 |
| Acetonitrile; trifluoroacetic acid (Bruker standard solvent) | Sigma-Aldrich | Cat# 900666 |
| α-D-Glucose | Sigma-Aldrich | Cat# 158968 |
| Guanidinoacetic acid | Novachem Pty Ltd | Cat# 070-00801 |
| Taurine | Sigma-Aldrich | Cat# 8086160005 |
| L-Glutamine | Sigma-Aldrich | Cat# G8540-10MG |
| L-Glutathione oxidized | Sigma-Aldrich | Cat# G4501-1G |
| L-α-Glycerophosphorylcholine | Sigma-Aldrich | Cat# G5291-10MG |
| L-α-Hydroxyglutaric acid | Sapphire Bioscience Pty Ltd | Cat# 21123-10MG |
| Digitonin | Promega | Cat# G9441 |
| IGEPAL CA-630 | Sigma-Aldrich | Cat# I8896-50ML |
| Bovine Serum Albumin (BSA) | Scientifix | Cat# BSASAU-0.1 |
| Trypan Blue | Bio-Rad | Cat# 1450013 |
| Magnesium chloride | Sigma-Aldrich | Cat# M8266-1KG |
| Sodium chloride | Sigma-Aldrich | Cat# 71380-500G |
| Tris-HCl | Promega | Cat# H5123 |
| RNase Inhibitor | Millipore Sigma | Cat# 3335399001 |
| DTT | Millipore Sigma | Cat# 646563 |
| Low TE Buffer (10 mM Tris-HCl pH 8.0, 0.1 mM EDTA) | Thermo Fisher Scientific | Cat# 12090-015 |
| SPRIselect Reagent Kit | Beckman Coulter | Cat# B23318 |
| Glycerin (glycerol), 50% (v/v) Aqueous Solution | Ricca Chemical Company | Cat# 3290-32 |
| Qiagen Buffer EB | Qiagen | Cat# 19086 |
| Isopentane (2-methylbutane) | Merck Life Science Pty Ltd | Cat# 277258 |
| Sodium dodecyl sulfate (SDS) | Merck Life Science Pty Ltd | Cat# L3771 |
| Hematoxylin | Merck Life Science Pty Ltd | Cat# MHS16 |
| Eosin | Merck Life Science Pty Ltd | Cat# HT110116 |
| **Critical commercial assays** |  |  |
| Xenium Slides & Sample Prep Reagents | 10x Genomics | Cat# PN-1000460 |
| Xenium Decoding Consumables | 10x Genomics | Cat# PN-1000487 |
| Xenium Instrument Accessory Kit, Module A | 10x Genomics | Cat# PN-1000530 |
| Xenium custom gene expression panels (panel A: 339 genes, panel B: 338 genes) | 10x Genomics | Panel A: ZQ94XW<br>Panel B: PAGXZA |
| Chromium Next GEM Single Cell Multiome ATAC + Gene Expression | 10x Genomics | Cat# PN-1000285 |
| Nuclei Isolation Kit, Nuclei PURE Prep | Sigma-Aldrich | Cat# NUC201 |
| ITO-coated glass slides (70-100 Ω) | SPI Supplies | Cat# 06400-AB |
| ACD RNAscope™ HiPlex Pro for COMET™ | Advanced Cell Diagnostics | ADV323680 |
| **Deposited data** |  |  |
| Xenium spatial transcriptomics, spatial metabolomics and immunofluorescence images | This paper | BioImage Archive: |
| single-nuclei RNA-seq and single-nuclei ATAC-seq multiome data | This paper | N/A |
| SMINT spatial transcriptomics and metabolomics data | Kriel et al.<sup>11</sup> | BioImage Archive: S-BIAD1426 |
| Safusidenib trial snRNA-seq data | Drummond et al.<sup>7</sup> | phs003976 |
| Flex scRNA-seq data for the reference atlas | Drummond et al.<sup>7</sup> | GSE292732 |
| Public WHO grade 2 and 3 mIDH glioma single-cell datasets for the reference atlas (and held-out grade WHO 4 tumors) | Johnson et al.<sup>2</sup>;<br>Tang et al.<sup>7</sup>;<br>Vallentgoed et al.<sup>31</sup>;<br>Spitzer et al.<sup>32</sup>; | GSE326221, GSE260928, CCell_4083 |
| Public bulk expression data used for 1p/19q validation | TCGA-LGG | https://portal.gdc.cancer.gov |
| Human Metabolome Database 5.0 | Wishart et al. | https://hmdb.ca/ |
| RaMP 3.0.7 | Braisted et al. | https://rampdb.nih.gov/ |
| GRCh38-2024-A human reference genome | 10x Genomics | https://www.10xgenomics.com/support/software/cell-ranger-arc/downloads |
| Ivy Glioblastoma Atlas Project (Ivy GAP) anatomic region classification | Puchalski et al. | https://glioblastoma.alleninstitute.org/ |
| Human prefrontal cortex single-cell data, gestation to adulthood | Herring et al. | https://doi.org/10.1016/j.cell.2022.09.039 |
| OPC-GABA cell state signature | Published study | https://doi.org/10.1016/j.ccell.2024.08.009 |
| **Experimental models: Organisms/strains** |  |  |
| Mouse: NPAI glioma model (NRAS-G12V, IDH1-R132H, p53 and Atrx loss) | Núñez et al. | https://doi.org/10.1126/scitranslmed.aaq1427 |
| **Oligonucleotides** |  |  |
| RNAscope™ HiPlex CS Probe – Hs-CLVS1-T1 | Advanced Cell Diagnostics | Cat# ADV1154487T1 |
| RNAscope™ HiPlex CS Probe – Hs-MME-T2 | Advanced Cell Diagnostics | Cat# ADV410897T2 |
| RNAscope™ HiPlex CS Probe – Hs-KCNH7-T3 | Advanced Cell Diagnostics | Cat# ADV1115427T3 |
| RNAscope™ HiPlex CS Probe – Hs-RIT2-T6 | Advanced Cell Diagnostics | Cat# ADV534357T6 |
| RNAscope™ HiPlex CS Probe – Hs-CAMK4-T7 | Advanced Cell Diagnostics | Cat# ADV1609447T7 |
| RNAscope™ HiPlex CS Probe – Hs-PDGFRA-T9 | Advanced Cell Diagnostics | Cat# ADV604487T9 |
| RNAscope™ HiPlex CS Probe – Hs-SPARCL1-T10 | Advanced Cell Diagnostics | Cat# ADV587527T10 |
| RNAscope™ HiPlex CS Probe – Hs-DGKG-T11 | Advanced Cell Diagnostics | Cat# ADV536967T11 |
| **Software and algorithms** |  |  |
| R 4.2.3; 4.5.3; 4.6.1 | R Core Team | https://www.r-project.org/ |
| Python 3.11 | Python Software Foundation | https://www.python.org/ |
| SMINT 2.0.0 | Kriel et al.<sup>11</sup> | https://zenodo.org/records/23684022 |
| SpaMTP 0.99.9 | Causer et al.<sup>12</sup> | https://github.com/GenomicsMachineLearning/SpaMTP |
| Xenium Onboard Analysis 2.0 | 10x Genomics | https://www.10xgenomics.com/support/software/xenium-ranger/latest |
| Xenium Ranger 4.0.0 | 10x Genomics | https://www.10xgenomics.com/support/software/xenium-ranger/latest |
| Xenium Explorer 3.0 | 10x Genomics | https://www.10xgenomics.com/support/software/xenium-explorer/latest |
| Zen Blue 3.8 | Zeiss | N/A |
| FIJI/ImageJ 1.54f | Schindelin et al. | https://imagej.net/software/fiji/ |
| Cellpose 2.0 | Stringer et al. | https://github.com/MouseLand/cellpose |
| bioformats2raw 0.9.4; raw2ometiff | Glencoe Software | https://github.com/glencoesoftware |
| dask and dask distributed 2025.7.0 | dask.org | https://github.com/dask/dask |
| zarr-python 3.1.1 | Miles et al. | https://zenodo.org/records/3773450 |
| scikit-image 0.25.2 | van der Walt et al. | https://doi.org/10.7717/peerj.453 |
| sf 1.0-19; lwgeom 0.2-14 | Pebesma and Bivand | https://CRAN.R-project.org/package=sf |
| sfheaders 0.4.4 | Cooley | https://CRAN.R-project.org/package=sfheaders |
| STalign 1.0 | Clifton et al. | https://github.com/JEFworks-Lab/STalign |
| SpatialData 0.2.3 | Marconato et al. | https://github.com/scverse/spatialdata |
| napari 0.5.4 | Sofroniew et al. | https://napari.org/ |
| Cardinal 3.0.1 | Bemis et al. | https://bioconductor.org/packages/Cardinal |
| SciPy 1.14.1 | Virtanen et al. | https://pypi.org/project/scipy/ |
| Cell Ranger ARC 2.0.2 | 10x Genomics | https://www.10xgenomics.com/support/software/cell-ranger-arc/latest |
| CellBender 0.3.2 | Fleming et al. | https://github.com/broadinstitute/CellBender |
| Scrublet 0.2.3 | Wolock et al. | https://github.com/swolock/scrublet |
| scater 1.34.1 | McCarthy et al. | https://github.com/jimhester/scater |
| scran 1.34.0 | Lun et al. | https://github.com/MarioniLab/scran |
| scuttle 1.16.0 | McCarthy et al. | https://github.com/LTLA/scuttle |
| escape 2.2.3 | Borcherding et al. | https://github.com/BorchLab/escape |
| SingleR 2.6.0; 2.8.0 | Aran et al. | https://github.com/dviraran/singler |
| SingleCellExperiment 1.28.1 | Amezquita et al., 2020 | https://bioconductor.org/packages/SingleCellExperiment |
| msigdbr 26.1.0 | Dolgalev, CRAN | https://cran.r-project.org/package=msigdbr |
| bluster 1.16.0 | Lun et al. | https://github.com/LTLA/bluster |
| limma 3.62.2; 3.66 | Ritchie et al., 2015; Wu & Smyth, 2012; Law et al., 2014 | https://bioconductor.org/packages/limma |
| edgeR 4.4.2 | Robinson et al., 2010 | https://bioconductor.org/packages/edgeR |
| MACS3 3.0.1 | Doherty et al. | https://github.com/macs3-project/MACS |
| Harmony 2.0.5 | Korsunsky et al. | https://github.com/immunogenomics/harmony |
| ggtern 4.0.0 | Hamilton et al. | https://github.com/nicholasehamilton/ggtern |
| Integrated Genomics Viewer 3.0.0 (IGV) | Robinson et al. | https://igv.org/ |
| Seurat 5.1.0 | Hao et al. | https://CRAN.R-project.org/package=Seurat |
| scvi-tools (scVI) 1.1.6 | Lopez et al.; Gayoso et al. | https://scvi-tools.org/ |
| CytoSPACE 1.1.0 | Vahid et al. | https://github.com/digitalcytometry/cytospace |
| GraphSAGE, StellarGraph 1.2.1 implementation | Hamilton et al. | https://github.com/williamleif/GraphSAGE |
| tidymodels 1.4.1 | Kuhn et al. | https://www.tidymodels.org |
| NeighbourNet 1.0.0 | Deng et al. | https://github.com/meiosis97/NeighbourNet |
| ArchR 1.0.3 | Granja et al. | https://github.com/GreenleafLab/ArchR |
| fgsea 1.32.4 | Korotkevich et al. | https://github.com/alserglab/fgsea |
| Immunomodulatory program framework | 10.1038/s41586-025-08633-8 | https://github.com/BernsteinLab/Calculate_Myeloid_cNMF_Usage/ |
| Panel calibration of immunomodulatory programs; T-ROS classifier; analysis code | This paper |  |
| Lunaphore Control Software v1.2.1.0 | Advanced Cell Diagnostics | https://lunaphore.com/products/comet/ |
| Bioconductor 3.22 | Huber et al., 2015 | https://bioconductor.org/ |
| org.Hs.eg.db 3.22.0 | Carlson, Bioconductor | https://bioconductor.org/packages/org.Hs.eg.db |
| AnnotationDbi 1.72.0 | Pagès et al., Bioconductor | https://bioconductor.org/packages/AnnotationDbi |
| SummarizedExperiment 1.40.0 | Morgan et al., Bioconductor | https://bioconductor.org/packages/SummarizedExperiment |
| ggplot2 4.0.3 | Wickham, 2016 | https://ggplot2.tidyverse.org/ |
| dplyr 1.2.1 | Wickham et al., 2019 | https://dplyr.tidyverse.org/ |
| tidyr 1.3.2 | Wickham et al., 2019 | https://tidyr.tidyverse.org/ |
| readr 2.2.0 | Wickham et al., 2019 | https://readr.tidyverse.org/ |
| forcats 1.0.1 | Wickham et al., 2019 | https://forcats.tidyverse.org/ |
| stringr 1.6.0 | Wickham et al., 2019 | https://stringr.tidyverse.org/ |
| knitr 1.51 | Xie, CRAN | https://yihui.org/knitr/ |
| rmarkdown 2.31 | Allaire et al., CRAN | https://rmarkdown.rstudio.com/ |
| ggrepel 0.9.8 | Slowikowski, CRAN | https://cran.r-project.org/package=ggrepel |
| scales 1.4.0 | Wickham et al., CRAN | https://cran.r-project.org/package=scales |
| svglite 2.2.2 | Wickham et al., CRAN | https://cran.r-project.org/package=svglite |
| igraph 2.3.4 | Csárdi & Nepusz, 2006 | https://cran.r-project.org/package=igraph |
| ggraph 2.2.2 | Pedersen, CRAN | https://cran.r-project.org/package=ggraph |
| tidygraph 1.3.1 | Pedersen, CRAN | https://cran.r-project.org/package=tidygraph |
| PyCave 3.2.1 | Borchert | https://github.com/borchero/pycave |
| MSigDB 26.1.1 | Subramanian et al., 2005; Liberzon et al., 2015 | https://www.gsea-msigdb.org/ |
| LabSolutions 5.1.28 (iMScope QT) | Shimadzu | https://www.ssi.shimadzu.com/products/liquid-chromatograph-mass-spectrometry/lc-ms-software/labsolutions-lcms/index.html |
| Imaging MS Solution 2.02 | Shimadzu | https://www.ssi.shimadzu.com/products/life-science-lab-instruments/imaging/imscope-qt/index.html |
| iMLayer Touch Panel software 26359023_E_12010 | Shimadzu | https://www.ssi.shimadzu.com/products/life-science-lab-instruments/imaging/imlayer/index.html |
| iMLayer Control Board software 26359022_5 | Shimadzu | https://www.ssi.shimadzu.com/products/life-science-lab-instruments/imaging/imlayer/index.html |
| iMLayer AERO 1.02.00 | Shimadzu | https://www.ssi.shimadzu.com/products/life-science-lab-instruments/imaging/imlayer-aero/index.html |
| hexbin 1.28.3 | Carr et al., CRAN | https://CRAN.R-project.org/package=hexbin |
| interp 1.1-6 | Gebhardt et al., CRAN | https://CRAN.R-project.org/package=interp |
| SpaMTPdb 0.99.4 | Causer et al. | https://github.com/GenomicsMachineLearning/SpaMTP |
| AUCell (via escape 2.2.3) | Aibar et al., 2017 | https://bioconductor.org/packages/AUCell |
| **Other** |  |  |
| Xenium Analyzer | 10x Genomics | N/A |
| Chromium Controller | 10x Genomics | N/A |
| Zeiss LSM 980 confocal microscope | Zeiss | N/A |
| Plan-Apochromat 20x/0.8 NA objective | Zeiss | Cat# 4406409903 |
| Plan-Apochromat 10x/0.45 NA objective | Zeiss | Cat# 4206419910 |
| CryoStar NX70 cryostat | Thermo Fisher Scientific | N/A |
| Leica AutoStainer ST5010 | Leica Biosystems | N/A |
| iMScope QT MALDI-TOF imaging mass spectrometer | Shimadzu | **N/A** |
| iMLayer and iMLayer AERO matrix deposition systems | Shimadzu | **N/A** |
| Alpha 3-4 LSCbasic freeze dryer | CHRIST | Version:<br>Controller: 1.5.4.1 (0)<br>IO Module 0x60: 1.0.2.0 |
| COMET | Lunaphore | N/A |
| NovaSeq X Plus sequencer | Illumina | N/A |

### EXPERIMENTAL MODEL AND STUDY PARTICIPANT DETAILS

#### Experimental model details

All animal experiments presented in this study were conducted according to regulatory standards approved by the Walter and Eliza Hall Institute Animal Ethics Committee (WEHI AEC 2024.002).

Pure C57BL/6 littermates were intracranially injected with 50,000 NPAI (*NRAS*-G12V and *IDH1*-R132H mutant, with *p53* and *Atrx* loss) tumor cells^57^ in 2 µL PBS and housed for the latency of the experiment under a 12-hour light/dark cycle. All animals were used in scientific experiments for the first time. This includes no previous exposures to pharmacological substances. Health status was normal for all animals.

#### Study participant details

Fresh tumor and Formalin Fixed Paraffin Embedded (FFPE) tissue was obtained from consented patients via the Royal Melbourne Hospital Neurosurgery Unit and the Royal Melbourne Hospital Neurosurgery Brain and Spine Tumour Tissue Bank (Melbourne Health Ethics #2020.214). Samples were de-identified and analyzed under a protocol approved by the Walter and Eliza Hall Institute Human Research Ethics Committee (HREC 21/21) and the Royal Melbourne Hospital Human Research Ethics Committee (HREC 2022.022) (**Table S10**). Age and sex were collected for all patients (**Figure S1A**) but were not used, unless stated, as variables in downstream analysis.

### METHOD DETAILS

#### Experimental data acquisition

##### Sample preparation

Tissue for deeply profiled SM/ST serial sections were flash-frozen immediately after resection in isopentane cooled in a liquid nitrogen bath to preserve endogenous metabolites, and stored at −80 °C until sectioning. Frozen tissue was sectioned without OCT embedding medium on a CryoStar NX70 cryostat at −20 °C. Blocks were mounted, equilibrated to chamber temperature and trimmed to the region of interest. Slides and samples were kept cold throughout sectioning to minimize freeze–thaw cycles. To capture intra-tumoral heterogeneity, each block was sampled across ∼480 µm using up to eight serial levels spaced 60 µm apart. At each level, three adjacent 10 µm sections were collected: histopathology (hematoxylin and eosin (H&E)), Xenium slide (10X-1000460, 10x Genomics) for ST followed by IF, and indium tin oxide (ITO)-coated slide for SM (**Figure S1A**).

##### H&E staining and histopathology

H&E sections were thawed, formalin-fixed and stained on a Leica AutoStainer (ST5010) and were also used to assess section quality. Slides were reviewed by certified neuropathologists (S.R.T, O.E.F.) and annotated according to the Ivy GAP classification^52^.

##### 10X Xenium ST

ST was performed with the Xenium platform (10x Genomics) using custom 338-gene and 339-gene panels with 307 overlapping genes (**Table S1**). As described previously, the protocol followed our previous work and the manufacturer’s fresh-frozen workflow^23^. Sections (10 µm) on Xenium slides were fixed in 4 % paraformaldehyde for 30 min at room temperature, then permeabilized with a 1 min wash in 1 X PBS, a 2 min incubation in 1 % SDS and further 1min washes in 1 X PBS. Slides were incubated in pre-chilled 70 % methanol for 60 min on ice and washed twice in 1 X PBS for 1 min each.

Probes were hybridized to target RNA overnight at 50 °C using 34 µL of probes per slide in Xenium Probe Hybridization Buffer (10 Genomics, PN-2000390) and TE buffer. Unbound probes were removed with PBS-T (1 X PBS, 0.05 % Tween 20). Ligation was performed with 481.2 µL Xenium Ligation Buffer (10x Genomics, PN-2000391), 13.8 µL Xenium Ligation Enzyme A (10x Genomics, PN-2000397) and 55.0 µL Xenium Ligation Enzyme B (10x Genomics, PN-2000398) per reaction, and circularized probes were enzymatically amplified with 495.0 µL Xenium Amplification Mix (10x Genomics, PN-2000392) and 55.0 µL Xenium Amplification Enzyme (10x Genomics, PN-2000399) per reaction. Autofluorescence was quenched with 500 µL of Xenium Autofluorescence Mix (10x Genomics, PN-2000753) in 100 % ethanol, and nuclei were stained with 500 µL Xenium Nuclei Staining Buffer (10x Genomics, PN-2000762). Slides were then loaded onto the Xenium Analyzer (version 2.0.0) for imaging and decoding. Raw outputs were processed with Xenium Ranger (version 4.0.0).

##### Immunofluorescence and imaging

###### Post-Xenium immunofluorescence

After Xenium acquisition, the same slides were immunostained as described previously^23^; no additional fixation or permeabilization was required. Slides were incubated in blocking buffer (0.1 % Tween 20 in 1 X PBS with 5 % goat serum) for 1 h at 4 °C and washed three times with PBS-T. Primary antibodies against GFAP (Rabbit monoclonal anti-GFAP; 1:1000) and MAP2A (Chicken polyclonal anti-MAP2A, PA1-10005, 1:1000) were diluted in blocking buffer and applied overnight at 4 °C. Slides were washed three times with cold PBS-T and incubated for 1 h at room temperature with secondary antibodies diluted 1:1000 in blocking buffer (goat anti-mouse Alexa 594; goat anti chicken-Alexa Fluor 488) together with Alexa Fluor 647-conjugated phalloidin to label filamentous actin 1:1000. After three further washes in PBS-T, nuclei were counterstained with DAPI (1:1000, 10 min, room temperature). Slides were washed three times in PBS-T, mounted in Fluoromount and stored at 4 °C until imaging.

###### Confocal image acquisition

Slides were imaged on a Zeiss LSM 980 confocal microscope. To ensure that the stage recognized the Xenium slide as a conventional 76 × 26 mm slide, support points were assigned to each corner of the Xenium slide sample area for stage calibration, and the Zen Blue experiment file was reused for subsequent imaging rounds. A low-resolution transmitted-light (T-PMT) tile scan (256×256 pixels) of the whole sample area was acquired with a Plan-Apochromat 10 x/ 0.45 NA objective, and regions of interest were drawn for each section. Depending on section size, 10 – 20 support points were placed per section so that all tiles were captured in focus. Using a Plan-Apochromat 20 x/ 0.8 NA objective, laser power was set for the 405, 488, 594 and 647 nm lines, and the z-position of each support point was calibrated manually to a plane in the middle of the section. Tiles of 3852 × 3852 pixels were acquired at LSM scan speed 7 in bidirectional mode, with z-stacks of 5 – 6 slices (z-step) centered on the calibrated focal plane.

###### Image processing and export

Maximum-intensity orthogonal projections were generated in Zen Blue (version 3.8). Tiles were exported as TIFF files and the projections stitched in Zen Blue with 10 % tile overlap. Whole-slide images were converted to OME-Zarr/OME-TIFF with bioformats2raw (version 0.9.4) and raw2ometiff for distributed processing and visualization.

##### RNAscope and sequential immunofluorescence

Multiplexed RNA detection was performed on the Lunaphore COMET platform using the RNAscope HiPlex Pro for COMET kit (Advanced Cell Diagnostics, ADV323680). Eight target probes were hybridized and detected sequentially across three rounds, with fluorophore assignment by tail: CLVS1 (Cat# 1154487; probe 1154487-T1, FITC), MME (Cat# 410897; 410897-T2, TRITC), KCNH7 (Cat# 1115427; 1115427-T3, Cy5), RIT2 (Cat# 534357; 534357-T6, TRITC), CAMK4 (Cat# 1609447; 1609447-T7, Cy5), PDGFRA (Cat# 604487; 604487-T9, FITC), SPARCL1 (Cat# 587527; 587527-T10, TRITC) and DGKG (Cat# 536967; 536967-T11, Cy5). Probes were detected in round 1 (CLVS1, MME and KCNH7), round 2 (RIT2 and CAMK4) and round 3 (PDGFRA, SPARCL1 and DGKG), with nuclei counterstained with DAPI (1:1000) in each round. After RNAscope detection, the same sections underwent sequential immunofluorescence (seqIF) on COMET, with three automated cycles of primary and secondary antibody incubation, imaging and antibody elution. Primary antibodies were incubated for 4 min and detected with goat anti-mouse 555 (TRITC channel) and goat anti-rabbit 647 (Cy5 channel) secondary antibodies (Lunaphore). Cycle 1 comprised mouse anti-GFAP (clone GA5; Cell Signaling Technology 3670; 1:700) and rabbit anti-OLIG2 (clone E6G6Q; Cell Signaling Technology 65915S; 1:200), cycle 2 rabbit anti-MAP2 (clone EPR19691; Abcam ab183830; 1:5000) and cycle 3 rabbit polyclonal anti-CLVS1 (Thermo Fisher Scientific OSC00321W-100UL; 1:5000), all reactive with human tissue. Nuclei were counterstained with DAPI, and autofluorescence and background were imaged in each cycle and subtracted Images were processed in Python (version 3.11.11) using tifffile (version 2026.3.3), NumPy (version 2.4.5), SciPy (version 1.17.1), scikit-image (version 0.26.0) and pandas (version 2.2.3). Residual offsets between imaging cycles were measured by band-pass filtering each channel and cross-correlating it with the protein autofluorescence image from the same filter set, in up to seven tissue regions (560 × 560 µm). The median integer-pixel shift across regions with reliable correlation was applied to each channel. Channels whose signal obscured the shared tissue texture took their shift from the elution image of the same cycle or from another channel in that cycle. Background was then subtracted pixelwise, with negative values set to zero. RNAscope channels were corrected with the matched autofluorescence image acquired at the RNA detection exposure. Protein channels were corrected with the pre-antibody autofluorescence image or where off-target secondary antibody signal was present. Nuclei were segmented in per section (560 × 560 µm) from Gaussian-smoothed DAPI (σ = 2 px) by thresholding, morphological opening and distance-transform watershed. Cell areas were approximated by expanding each nucleus by ∼3 µm. Olig2 was scored as mean nuclear intensity. GFAP and MAP2A were scored as mean intensity in the perinuclear ring (cell area minus nucleus), relative to the local channel mean (Gaussian σ = 180 px, ∼50 µm), so that cells were called positive only if they stood out from the surrounding fibre network. Cells were called Olig2+ at a nuclear intensity above 80, GFAP+ at a relative intensity above 1.15, and MAP2A+ at a relative intensity above 1.30 together with an absolute ring intensity above 600. Cells were then assigned, in order, as MAP2A+ (Olig2-negative), Olig2+GFAP+, Olig2+ or GFAP+; remaining cells were marker-negative.

RNAscope transcripts were detected as local maxima of a white top-hat-filtered image (Gaussian σ = 1 px, 9 px structuring element), and assigned to the cell containing them. Per-probe thresholds were set at the median + 8 robust SD (1.4826 × MAD) of the top-hat signal of the matched autofluorescence image within tissue. A probe was considered detected in a cell at ≥ 2 transcripts, and cells with ≥ 6 of the 8 probes (*CLVS1, MME, KCNH7, RIT2, CAMK4, PDGFRA, SPARCL1, DGKG*) were defined as program-high.

##### MALDI-TOF SM

Cryosections were mounted onto indium tin oxide (ITO)–coated slides (70–100 Ω), freeze-dried for 30 min and optically imaged before matrix deposition. An iMLayer AERO sprayer (Shimadzu) applied eight layers of α-cyano-4-hydroxycinnamic acid (CHCA; 10 mg/mL in acetonitrile/water, 50:50 v/v, containing 0.1 % trifluoroacetic acid), with a stage speed of 70 mm/s, 1 s drying intervals and a 5 cm nozzle distance. MALDI-TOF SM was performed using an iMScope QT instrument (Shimadzu) at 10 µm spatial resolution. Acquisition settings comprised a laser intensity of 60, detector voltage of 2.36 kV, laser repetition frequency of 2,000 Hz and desolvation line temperature of 250 °C, with 50 laser shots accumulated per pixel, as described previously^24^.

##### 10X snRNA and ATAC-seq and sequencing

Tumor tissue was frozen on dry ice and stored at -80 °C until nuclei isolation. All steps involved in isolation of nuclei were conducted using pre-chilled buffers and a centrifuge pre-set to 4 °C. Tissue was transferred to a Dounce homogenizer containing lysis buffer (10 mM Tris-HCl pH 7.5, 3 mM Magnesium chloride (MgCl_2_), 10 mM Sodium chloride (NaCl), and 0.005 % IGEPAL CA-630) and incubated for 1 min before homogenization. Tissue was homogenized using a combination of loose and/or tight pestles and a 1 ml pipette tip, depending on the individual piece of tissue and incubated for 1 min. The homogenates were filtered through a 40 μm strainer and centrifuged at 500 × g for 5 mins. Supernatant was discarded, the pellet was resuspended in additional lysis buffer and 0.01 % Digitonin was added. Following an incubation of 3 mins, wash buffer (Dulbecco’s PBS containing 1 % bovine serum albumin (BSA)) was added to neutralize the reaction. An addition centrifugation step was performed at 500 × g for 5 mins, the supernatant was discarded and the pellet resuspended in fresh wash buffer. Next, nuclei were mixed with 1.8 M sucrose solution (prepared from 2 M Sucrose Cushion solution (S9308-475ML), Nuclei PURE Sucrose Cushion Buffer (S9058-120ML); Nuclei Isolation Kit, Nuclei PURE Prep, Sigma NUC201) and layered onto 1.3 M sucrose solution. The sucrose gradient was subsequently centrifuged at 3000 × g for 10 mins. The debris ring was wiped away with a Kimwipe and supernatant was discarded. Pellet was resuspended in a volume of Chromium Next GEM Single Cell Multiome Diluted Nuclei Buffer (PN-2000207) depending on pellet size. Nuclei were counted using trypan blue and a haemocytometer. For single-nuclei ATAC and gene expression sequencing of fresh nuclei, 10,000 nuclei per sample were targeted on the Chromium controller, after which library preparation was conducted using the Chromium Next GEM Single Cell Multiome ATAC + Gene Expression kit (10x Genomics), following the manufacturer’s instructions. Libraries were sequenced on a NovaSeqX Plus (Illumina) in paired-end mode, targeting 20,000 reads per nucleus for the RNA-sequencing libraries and 25,000 reads per nucleus for the ATAC-sequencing libraries. For the cohort, we found on average 7,103 reads per nuclei and 7,450 fragments per nuclei.

#### Alignment of spatial modalities

Adjacent sections from each level (H&E, Xenium/IF and spatial metabolomics) were brought into a common coordinate framework with SMINT^23^. Briefly, cell centroids were rasterized into density images with STalign^53^ (version 1.0), pre-aligned by rotation, and registered by large deformation diffeomorphic metric mapping (STalign.LDDMM) using manually placed landmarks, which jointly estimates an affine transformation and a diffeomorphism. The resulting transformation was applied to the source coordinates with STalign.transform_points_source_to_target.

#### Analysis of ST

##### Processing, filtering, normalization and clustering

ST data were preprocessed separately before integration. From raw transcript outputs from Xenium, transcripts were filtered to retain those with a distance of 0 to segmented nuclei (corresponding to transcripts within the nucleus) and high quality (qv > 20). Further filtering was performed to remove nuclei with ≥ 3 negative control probes, and nuclei with ≤ 20 transcripts or ≤ 5 unique genes. Filtered counts were log-normalized and clustered via PCA based on the first 25 principal components.

##### Annotation

Annotations were performed separately across samples enlisting semi-supervised annotation followed by manual refinement^11^. Individual cell annotations were estimated using SingleR (version 2.6.0) and two references^33,54^ to identify non-malignant populations and to characterize malignant characteristics. A Uniform Manifold Approximation and Projection (UMAP) projection was used to visualize the cell populations based on the first 25 principal components with default parameters, excluding min_dist, which was set to 0.1. Each cluster was preliminarily assigned to the most common annotation within the cluster. Clusters were then confirmed or modified based on overexpression of marker genes. These annotations were performed cyclically, first to define larger populations (Malignant: *BCAN, PTPRZ1*; Immune *RNASET2, ITGAX, PTPRC*: Vasculature: *IGFBP7, SPARC*; Glial: *APOD, SOX10*; Neuronal: *STMN2, NPTXR*), then reperforming clustering on these populations separately to pull out more specific annotations (AC-like: *NTRK2, PMP2*; Astrocyte: *AQP4, CST3*; B cell: *BANK1, CD79A*; *CD4*+ T cell: *IL7R, CD4*; *CD8*+ T cell: *CD8A, CCL5*; Dendritic cell: *TGFBI, HLA-DRB5*; Endothelial: *PECAM1, IGFBP3*; Excitatory Neuron: *EPHA4, LMO4*; Fibroblast: *FBLN1, APOD*; Inhibitory neuron: *SPOCK3*; Macrophage: *CD163, MS4A6A*; Microglia: *CX3CR1, P2RY12; GPNMB+* microglia / macrophage: *GPNMB, LPL*; *IL1B*+ microglia: *IL1B, CD83*; MES-like: *CHI3L1, IDO2*; Monocyte: *CTSS, COTL1*; Mural: *IGFBP7, SPARC*; NK cell: *KLRC1, CD247*; NPC-like: *MAP2, PROX1, TUBB2B*; Neutrophil: *ITGAX, CXCR1*; OPC: *CA10, SOX10*; OPC-like: *DLL3, SOX4*; Oligodendrocyte: *MBP, ABCA2*; Progenitor: *MKI67, TOP2A*; T Reg cell: *CTLA4, FOXP3*).

##### Integration across ST samples

Two custom gene panels were used to profile different samples with n =338 and 339 genes each, and n = 307 overlapping genes (**Table S1**). For integration purposes, samples were subsetted to only those overlapping genes. scVI (scvi-tools, version 1.1.6) was employed to integrate samples together, using the sample identity as the batch variable and raw counts as the input. A model was trained across 100 epochs, resulting in a latent-representation of 10 variables for each nucleus. Post-integration, Leiden clustering and UMAP dimension reduction were reperformed, and sample annotation concordance was validated.

##### Niche analysis

GraphSAGE (v1; StellarGraph version 1.2.1) was performed on transcripts passing quality control filtering^17,55^. Briefly, to infer structures in the data, transcript-based niches were identified using all detected nuclei transcripts as nodes, ignoring cell segmentation, and edges between nodes drawn based on Euclidean distances. Sample-specific graphs of 10 cases (A1 to A9 and O1) were assembled then subsampled to reduce memory usage, before aggregation into a single joined graph for model training (4,586,495 nodes, 113,592,432 edges). Other parameters used for graph construction and training remained as previously reported^17^. We trained the model on the merged graph with 10 epochs and achieved a training accuracy of 0.82.

We then extracted the embedding representation for all transcript nodes across all 26 samples using the trained model. All sample nodes were then clustered (102,129,436 total nodes) using GaussianMixtureModel (k = 12) from PyCave (version 3.2.1). For each sample, transcripts were consolidated into hexbin plots using hexbin (version 1.28.3). A consistent bin width of 70 µm was used to ensure uniform bin sizes across samples. Each bin was distinctly labeled and color coded based on the most dominant node label among the contained transcripts. Bins with < 10 transcripts were excluded from the analysis. Cell assignment to niches was determined by calculating the nearest hexbin centroid via the st_nearest_feature function (sf, version 1.0-19). Cell type enrichment was calculated as the proportion of cells assigned to each transcript niche, normalized across both niches and cell types.

##### Niche characterization: Ivy GAP regions

Region outlines were aligned to GraphSAGE hexbin planes by previously mentioned STAlign strategies (**Alignment of spatial modalities**). Hexbin assignment to region was performed using the sf package and sfheaders (version 0.4.4) via the st_join function. Niche density was calculated as the mean number of processed cells present in a single hexbin. Cell centroids were used to assign cells to individual hexbins using the sf_join function in sfheaders. Niche prevalence was calculated as the proportion of samples with ≥ 40 hexbins of a given niche present. This cutoff was chosen to remove samples with extremely low and potentially misrepresentative niche allocation.

##### Pathway Analysis

For each sample, we integrated spatial transcriptomics data with corresponding snRNA-seq data using CytoSPACE (version 1.1.0). As CytoSPACE assigns profiles only from cells with matched annotations, spatial annotations were collapsed into simplified aggregates to accommodate for low cell type abundance in the paired single-nuclei data. These aggregates included ‘Macrophage’ (macrophages and M2/ *GPNMB*+ macrophages), ‘Microglia’ (microglia, dendritic cells, monocytes and neutrophils), and ‘Lymphoid’ (*CD4*+ T cells, *CD8*+ T cells, NK cells, B cells and T Reg cells).

Gene-set activity was tested with cameraPR in limma (version 3.66) on the moderated t-statistics from the cell-type-adjusted voom model (each niche vs the mean of the others). P-values were Benjamini-Hochberg ^56^ (BH)-adjusted across niches, and effect size is reported as the mean log fold change of set genes.

##### Niche neighbor analysis

Niche neighbors were determined via the the st_join function of sf. For each hexbin, a buffer of 75 µm was extended from its centroid via st_buffer ensuring that any adjacent hexbins are captured using st_join. Aggregating niche identities of all adjacent hexbins of a target niche across all samples identifies its niche neighbors. Niche neighbor proportions were then compared to overall niche proportions to assess overrepresentation using an enrichment-based statistical approach.

##### Treatment enrichment

Enrichment of niches in opposing treatment cohorts was calculated by an enrichment-based statistical approach comparing the number of hexbins of a given niche in each cohort to the relative proportions of each niche overall.

#### Analysis of SM

##### Spectral preprocessing and feature extraction using Cardinal

Profile-mode MALDI-TOF MSI spectra acquired from tumor tissue were processed using Cardinal (version 3.0.1 in R version 4.2.3). The m/z range of 70 – 400 was divided into consecutive, non-overlapping intervals of 10 m/z. Within each interval, spectra underwent totalion-current normalization, with intensities scaled such that their summed absolute intensity equaled the number of native spectral bins. Spectra were subsequently smoothed using a Savitzky–Golay filter with a third-order polynomial and a five-bin window. Baseline correction was performed using the local-minimum method (locmin, window = 5), with linear interpolation between local minima and negative corrected intensities set to 0. Detectable CHCA peaks, including [M+H]+, [M+K]+, [M+Na]+ were excluded from the analysis.

Peaks were detected using Cardinal’s median absolute deviation (MAD) with a signal-to-noise threshold of 6, a 5-bin window and a single noise-estimation block. Detected peaks were aligned to reference positions calculated as intensity-weighted m/z centroids from the mean peak-picked spectrum. Alignment tolerances were estimated separately for each interval from the median relative spacing of adjacent m/z bins, expressed in ppm and rounded to the nearest 0.5 ppm. Features detected in ≥ 1 % of spectra were retained.

The preprocessed spectra were then binned against the retained reference peak list using area-based quantification (type = “area”). Peak intensities were calculated by summing the corrected intensities between the flanking local minima, using the same automatic tolerance-estimation procedure for reference matching. Missing peak matches were assigned zero intensity. Interval-specific feature matrices were combined for downstream analysis.

##### Metabolite annotation

###### Untargeted metabolite annotation

Following Cardinal preprocessing, feature intensities, registered spatial coordinates and associated metadata were organized in SpatialExperiment objects using SpaMTP (version 0.99.9 in R version 4.6.1). Putative metabolite annotations were generated using SpaMTP::annotateSM() against the RaMP (version 3.0.7) chemical-property and precomputed structural-feature resources accessed through SpaMTPdb (version 0.99.4). Feature m/z values were searched using the general positive-ion adduct rules and a mass tolerance of 20 ppm, without a matrix-specific adduct restriction. The minimum annotation score was set to zero, and candidate identities, molecular formulae, adduct assignments, mass errors and annotation scores were retained. Unmatched features were preserved for downstream analysis. Cross-section feature clustering used a separate tolerance of 50 ppm.

###### Standard-guided ion assignment and raw spectral assessment

For the targeted raw-profile analysis, theoretical ion masses were calculated from molecular formulae using a curated set of positive-mode adducts, including [M+H]⁺, [M+NH₄]⁺, [M+Na]⁺, [M+K]⁺, [M−H+2Na]⁺, [M−H+Na+K]⁺ and [M−H+2K]⁺. Predefined dimer and doubly charged forms were additionally evaluated for selected compounds. Where standards were available, candidate selection was restricted to standard-supported adducts. Standard support required a mass match within 50 ppm, positive mean intensity, an intensity rank ≤ 50 and a specificity score ≥ 5. Specificity was calculated as the mean intensity in the corresponding standard divided by the maximum matched mean intensity among the other standards plus 1.

Strong raw spectral support required the theoretical m/z to fall within the measured full width at half maximum (FWHM) in ≥ 75% of sections and a median absolute apex mass error of ≤ 10ppm among sections with a detected peak. Moderate support required corresponding thresholds of ≥ 50% and ≤ 20ppm. Candidate ranking incorporated predefined adduct assignments, standard evidence, mass-support class, adduct plausibility and tissue detection frequency. Weak or unsupported assignments were flagged.

A separate peak-separability assessment required an apex-to-baseline ratio ≥ 3 that exceeded the corresponding ratios of ≥ 90 % of neighboring local maxima. Neighboring peaks were evaluated within the stored local spectrum, with their baselines estimated from flanking minima within 40 native bins. Support in this assessment required theoretical-mass coverage by the FWHM in ≥ 75% of sections and distinguishable peaks in ≥ 50 % of evaluable sections.

##### Normalization and peak filtering

Pixel-wise spectra were processed using Cardinal, including Savitzky–Golay smoothing (smoothSignal(method=“sgolay”), default settings) and baseline correction (reduceBaseline). Peaks were detected using peakPick aligned using peakAlign and filtered using peakFilter (freq.min = 0.01), with a minimum detection frequency of 1 % of pixels within each dataset. Cardinal-processed pixel intensities were TIC-normalized and multiplied by 10 . To compare metabolite abundances across niches and patients, TIC-normalized intensities were converted to z-scores and compared using non-parametric Wilcoxon tests.

##### Metabolite Annotation Confidence Levels

Metabolites classified as Annotation Confidence Level 1^28^ underwent an independent experiment whereby 10mg metabolite standard was dropped (2 µL) onto a glass ITO slide and onto a 10 µm section of flash frozen mouse brain tissue mounted on a glass ITO slide. Standards were evaluated by MALDI-TOF mass spectrometry (MS, **MALDI-TOF SM**) and candidate peaks were further evaluated by MS/MS. For all metabolites, the respective precursor m/z values were selected for MS/MS acquisition using a Q1 isolation window of 1 Da. Product ion spectra were acquired using a collision energy of 20, with a collision energy spread of ± 5. Peaks matching rational adducts and those identified in the experimental slides were annotated as Annotation Confidence Level 1 (Glucose, Glutamate, Glutathione and GAA). Metabolite peaks identified as enriched in the MS/MS or MS trace and present in the experimental dataset but were not classical predicted adducts were annotated as Level 2 (Taurine, Glutamine, Citrate and others). Metabolites that were not assessed by standards as above, but had rational adducts were classified as Level 3 and those m/z peaks that had a reproducible analytical signal but low confidence metabolite annotation were classified as Level 4 (**Table S2**). For all experiments, MALDI-TOF SM was performed in positive-ion mode using CHCA as the matrix replicating the same experimental conditions as for patient samples (**MALDI-TOF SM**).

##### Niche metabolite profiles

Within each tissue section, MSI pixels were assigned to the spatial bin it resides in. Ion intensities were summed within each bin and then aggregated into niche-level profiles using the corresponding niche-composition weights according to overlap of SM pixel and ST niche hexbin. Effective support was calculated as the niche-weighted number of assigned MSI pixels.

##### 2-HG analysis

###### Raw spectral extraction and adduct assessment

To assess 2HG-associated signals and substrate-dependent adduct responses, flash-frozen brain sections from one C57Bl/6 mouse bearing an intracranial NPAI (*NRAS*-G12V and *IDH1*-R132H mutant, with *p53* and *Atrx* loss) tumor^57^ were mounted on ITO-coated slides. Three acquisitions were analyzed, corresponding to 2HG standard concentrations of 1, 5 and 10 mg/mL, each with one spot on exposed glass, tumor tissue and non-tumor tissue. Ion intensities were extracted within ± 20 ppm of theoretical masses for protonated and sodium- and potassium-containing adducts of 2HG (C₅H₈O₅) and its dehydrated formula (C₅H₆O₄). CHCA sodium and potassium adducts were examined to contextualize substrate-dependent responses. Raw and total-ion-current (TIC)-normalized intensities were analyzed, and regional mean spectra were inspected for corresponding local peaks. Mean spot intensities were compared across substrates, with substrate-matched annuli extending 300 – 800 µm beyond spot boundaries used to assess local background.

###### Control of spot-associated contamination and candidate screening

Unspotted tissue analyses excluded tissue edges by 100 µm and standard spots with an additional 300 µm margin. Candidates required higher mean intensity in tumor than non-tumor tissue in every acquisition under both raw and TIC-normalized measurements, consistency across 100, 300 and 500 µm spot-exclusion margins, and positive signal in ≥ 1 % of unspotted tumor pixels per acquisition. Standard-associated enhancement required ≥ 2-fold signal relative to unspotted tissue of the same type and ≥ 1.5-fold signal relative to local background, under both intensity measures. The stringent subset met both criteria in all six tissue spots; the extended subset met each criterion in ≥ 4 spots, with the criteria counted separately.

###### Quantification and statistical analysis

Exploratory tumor enrichment was assessed using equally weighted means from 500 × 500 µm spatial blocks retaining ≥ 40 % of their nominal area, with ≥ 4 blocks per tissue region. One-sided studentized permutation tests used 9,999 whole-block label permutations. BH correction^56^ was applied across all 3,575 windows separately for each acquisition and intensity measure; retention required adjusted p < 0.05 in all six comparisons. Sensitivity analyses varied mass tolerance, spot radius, local-background subtraction, and spatial block size and alignment.

#### Construction of scRNA-seq atlas

##### Publicly available data and their integration

Publicly available datasets^6,7,17,31,32^ chosen to capture the cellular and molecular diversity of mIDH tumors were identified from previously published studies and corresponding data repositories (**Table S4**). We collated all normal and malignant cells and nuclei obtained from grade 2 and 3 samples. For each dataset, available sample-level and cell-level metadata were collected and harmonized into a common metadata framework. Where applicable, clinical and sample characteristics including tumor grade, diagnosis, treatment status, sample identity, recurrence status and dataset/source, were standardized across datasets using consistent terminology and categorical definitions.

For each dataset, initial quality control was performed independently to remove low-quality cells and nuclei marked by high mitochondrial reads and low UMI counts according to data specific quality control criteria. Following quality control, datasets were integrated using the variational autoencoder implemented in the scVI toolkit^58^ (version 1.1.6), with each sample identity specified as the batch key and using default parameters. The resulting X_scVI latent representation was used for downstream processing. A UMAP embedding was next generated from the scVI reduced dimensions with spread set to 3 and mindist set to 0.1 with 15 nearest neighbors (n_neighbors). Importantly, the low-dimensional shared latent space representation of nuclei were only used for data integration, visualization and cell type annotation and was not used for differential gene expression analyses. The integrated atlas comprised 577,916 cells and nuclei. Utilizing the same integration approach, we performed a separate scVI integration restricted to cells and nuclei classified as malignant. Sample identity was used as the batch key again using default parameters. The integrated malignant-only dataset comprised 345,386 cells and nuclei.

##### Annotation

Cells and nuclei from the mIDH atlas, comprising both malignant and normal cell populations, were clustered using the scVI reduced dimensions via Leiden clustering using default parameters implemented in bluster (version 1.16.0). Clusters were manually annotated based on existing annotations from the integrated in-house datasets and refined using marker gene expression. For a first pass, all cells and nuclei of in-house datasets were automatically annotated using SingleR (version 2.8.0) and two reference datasets^33,54^. The resulting initial annotation of the mIDH atlas identified the major normal cellular compartments, including vasculature, immune, neuron, oligodendrocyte, astrocyte, OPC and radial glia (RG) (Vasculature: *IGFBP7*, *COL4A2*; Immune: *PTPRC*, *CD68*; Neuron: *NPTXR*, STMN2; Oligodendrocyte: *MBP*, *ERMN*; Astrocyte: *NTRK2*, *MGST1*; OPC: *CNP*, *CSPG4*; RG: *PAX6*). The vasculature, immune and neuron compartments were subsequently subsetted and independently reclustered using the same dimensionality reduction and Leiden clustering approach. Subclusters were manually annotated based on established marker gene expression. Within the vasculature compartment, mural, endothelial and fibroblasts were identified (Mural: *DCN*; Endothelial: *COL4A2*, *PECAM1*; Fibroblasts: *FBLN1*, *COL1A2*). The neuron compartment was further resolved into excitatory and inhibitory Neurons (Excitatory Neuron: *NRN1*, *HOPX*; Inhibitory Neuron: *GRIK1*, *KIT*). The immune compartment comprised myeloid (*PTPRC*, *RNASET2*) and lymphoid (*TRAC, CD3*) populations.

A similar annotation strategy was applied to the malignant-only mIDH atlas to further resolve malignant tumor cell states. Malignant cells and nuclei were reclustered using Leiden clustering. Clusters were manually annotated based on marker gene expression, identifying AC-like, OPC-like, NPC-like, MES-like and Progenitor tumor cell states (AC-like: A*QP4*, *APOE*; OPC-like: *APOD*, *OLIG1/2*; NPC-like: *STMN2*, *DCX*; MES-like: *CHI3L1*, *ITGB1*; Progenitor: *TOP2A*). Two clusters comprising dying or poor-quality cells were identified featuring higher levels of mitochondrial reads and were consequently excluded from further analysis. Following this filtering, the final combined and processed mIDH atlas comprised 574,870 cells and nuclei, while the malignant-only mIDH atlas comprised 342,340 cells and nuclei. In addition to the canonical malignant glioma cell states, we identified an additional previously unrecognized transcriptional population that we termed “*RIT2*+ tumor cells”.

##### Myeloid granular state annotation

The single-nuclei atlas was subset to the immune populations, and lymphoid cells split from myeloid. 4,794 nuclei originally clustering with immune cells and annotated as ‘Immune’ did not integrate with other immune populations and featured elevated expression of both immune and malignant-associated genes (**Figure S5A**); consequently, they were filtered from further analysis as inconclusive annotations. A subsequent scVI integration was performed to correct batch effects in the myeloid data. This integration was performed with an additional ‘patient’ covariate employed to enhance alignment across studies. Canonical marker genes and Leiden clustering identified the refined myeloid cell type annotations (Macrophage: *CD163*, *MS4A6A*; Microglia: *CX3CR1*, *P2RY12*; *IL1B*+ microglia: *IL1B*, *CD83*; *GPNMB*+ microglia/macrophages: *GPNMB*, *LPL*; Dendritic cells: *TGFBI*, *CD74*; Monocyte: *CTSS*, *COTL1*; Neutrophil: *ITGAX*).

##### Identification of marker genes

To further validate the newly defined *RIT2*+ tumor cell state, we identified a distinct transcriptional signature associated with the population. This signature was subsequently used for independent validation of the tumor cell state across datasets. To guide identification of a transcriptional signature for this population, we first performed a differentially expressed gene (DEG) analysis using limma and edgeR (version 4.4.2), comparing the *RIT2*+ tumor cell population with the neighboring AC-like and OPC-like populations combined. Other malignant populations were excluded from this comparison to generate a more specific transcriptional signal associated with the *RIT2*+ tumor cell state. Counts were aggregated at the sample level to create pseudobulk profiles using scuttle (version 1.16.0). To ensure statistical robustness, only aggregated profiles with ≥ 10 cells/nuclei were retained. We performed DEG analysis using the limma workflow to model the effect of cell state on gene expression while accounting for sample-level variability. The design matrix included terms for cell state (*RIT2*+ tumor cells versus AC-like/OPC-like) and sample identity as covariates, with the cell state contrast representing the effect of interest. A total of 1907 DEGs were defined for the *RIT2*+ tumor cell state with significance was determined by a FDR threshold of < 0.05.

To further refine the transcriptional signature, we used a regularized logistic regression model in combination with manual inspection to identify genes that discriminated the *RIT2*+ tumor cell state from neighboring AC-like and OPC-like tumor populations. Cells/nuclei were assigned a binary response variable representing *RIT2*+ tumor cells versus non-*RIT2*+ tumor cells (AC-like/OPC-like) status, and only genes identified as differentially expressed in the preceding analysis were considered as candidate predictors. A lasso model was fitted. Model performance was assessed using ROC AUC and a confusion matrix, and per-cell predicted probabilities of *RIT2*+ tumor cell-state membership were generated for visualization on UMAP embedding. Model coefficients were extracted to generate a ranked list of 195 informative genes and further refined using manual inspection.

##### Comparison to other datasets

###### OPC-GABA comparison

The OPC-GABA cell type has recently been described as a distinct population in glioma^35^. To assess its relationship to the *RIT2*+ tumor cell state, we compared the published OPC-GABA cell state signature derived from Patch-seq and single-cell RNA-sequencing data of grade 2 and 3 mIDH tumors with the defined AC-like, OPC-like and *RIT2*+ tumor cells in our mIDH glioma atlas. Expression of the published signature genes across these populations was visualized using a dot plot to compare their transcriptional profiles.

###### Developmental trajectory analysis

To assess the developmental dynamics of genes from our *RIT2*+ tumor cell state signature, we utilized a publicly available dataset capturing a normal human prefrontal cortex from gestation through to adulthood^59^. We examined the temporal expression patterns of selected *RIT2*+ tumor cell state markers, *CAMK4*, *KCNH7*, *KIRREL3*, *RIT2* and *PTPRT*, across astrocytes and OPCs. Mean log-normalized expression was calculated for each gene within each cell type and timepoint, then scaled row-wise. Expression trajectories were ordered chronologically and smoothed using Locally Estimated Scatterplot Smoothing (LOESS) with a span of 1.5 to visualize broad temporal trends in gene expression.

To further assess the cell type specificity of *RIT2*+ tumor cell state marker *NELL1* within the normal developmental context, we compared average log-normalized expression across OPC, oligodendrocyte and astrocyte populations in the same dataset. Mean expression was calculated for each gene within each cell type and visualized as bar plots.

###### RIT2+ tumor cell state grade comparison

To independently assess the representation of the defined *RIT2*+ tumor cell population, we subsequently integrated held-out grade 4 mIDH glioma samples from the respective data sources that were excluded from the initial malignant-only atlas integration^6,7,31^. The same integration and annotation framework was applied, and the established tumor cell state definitions were used to annotate the integrated dataset.

###### TCGA gene expression analysis

We analyzed the TCGA low grade glioma cohort using bulk RNA-sequencing data to validate the presence of the *RIT2*+ tumor cell state transcriptional program in bulk tumor samples. Gene set enrichment scores were calculated using AUCell implemented through escape (version 2.2.3) for the *RIT2*+ tumor cell state signature, with a minimum gene set size of 10 genes. An *RIT2*+ tumor cell state signature score was extracted for each sample and merged with corresponding clinical and molecular metadata for downstream analysis. The resulting *RIT2*+ tumor cell state signature score was compared across *IDH*-codeleted molecular subtypes within the TCGA cohort.

##### GSEA

We conducted gene-set enrichment analysis (GSEA) from ranked lists generated using results from DEG analysis. For the *RIT2*+ tumor cells versus AC-like/OPC-like analysis, we created a ranked list of genes using −log10(p-value) while accounting for the direction of log-fold change. GSEA was then performed using fgsea (version 1.32.4), including a total of 10,520 gene sets in the analysis: 10,461 Gene Ontology gene sets and 50 hallmark gene sets obtained from mSigDB (version 26.1.1), as well as 4 malignant and 5 nonmalignant gene sets.

To assess GABAergic gene module expression, we evaluated signature expression across malignant and neuronal populations for each patient. A one-sided paired Wilcoxon signed-rank test was used to compare *RIT2*+ tumor cells to the equally weighted mean across other malignant populations, including populations with ≥ 20 cells per patient.

##### Gene regulatory network inference

To investigate the regulatory architecture governing the *RIT2*+ tumor cells, we applied NeighbourNet^36^ (version 1.0.0). Briefly, NeighbourNet estimates cell-specific gene co-expression by combining dimensionality reduction with local regression. Embedded gene expression is used to infer k-nearest neighbor (knn) relationships for each cell and local regression models subsequently estimate gene expression repeatedly with predictor contributions as a measure of weighted gene co-expression. As *RIT2*+ tumor cells were predominately associated with oligodendrogliomas, we first subsetted the mIDH atlas to this diagnosis before proportionally downsampling malignant cell states within each dataset source to a total maximum of 5000 cells/nuclei. Rather than using the default PCA space, the NeighbourNet nearest neighbor graph was constructed using the scVI latent representation of the integrated dataset to minimize potential batch effects between dataset sources. NeighbourNet was then applied to the proportionally downsampled dataset using default parameters, using publicly available receptor and gene lists provided by the package. The response gene list was curated to include canonical glioma gene sets. Top *RIT2*+ tumor cell state signature genes as well as genes associated with development and differentiation.

NeighbourNet identified 20 distinct meta-networks (MNs), which were visualized as a heatmap showing predicted regulatory weights across malignant cell states. MN3 was identified as a regulatory program uniquely representing *RIT2*+ tumor cells. To visualize the learned NeighbourNet MN3 space, cells were projected onto the first two MN coordinates and colored according to their annotated tumor cell state. The top predicted regulators and targets within MN3 were further partitioned into four modules using prepare.visualise with n.clu set to 4.

##### Immunomodulatory signature

Myeloid immunomodulatory programs were calculated using the consensus myeloid program calculator shiny app (https://github.com/BernsteinLab/Calculate_Myeloid_cNMF_Usage/), using log-normalized counts as the input, subsetted to only the genes within both the cNMF myeloid program signatures and the myeloid cells within the reference single cell dataset (n = 1,789). Results were visualized in a grid arrangement consistent with the original publication.

#### Analysis of snRNA- and ATAC-seq

##### Processing, filtering, normalization and integration

Data was first processed using Cell Ranger ARC (10X Genomics, version 2.0.2) to align ATAC and gene expression FASTQ files to the GRCh38-2024-A reference genome. Gene expression reads were quantified using unique molecular identifier (UMI) counting, while ATAC reads were used to generate chromatin accessibility fragments and peak counts.

###### snRNA-seq

CellBender (version 0.3.2) was applied to each sample using default parameters to estimate and remove ambient RNA. To identify and remove doublets, we employed scrublet (version 0.2.3), setting an estimated threshold based on the multiplet rate, with an upper limit cut-off of 0.25. We calculated mitochondrial expression percentage for nuclei using scater (version 1.34.1) and identified outlier nuclei using the MAD approach to dynamically set thresholds for minimum UMI counts and minimum detected genes. We filtered nuclei using the computed thresholds and removed those with a mitochondrial expression percentage > 10 %. Overall, 21,418 nuclei passed quality controls, with on average 2,655 genes detected per nuclei. We then applied a log-normalization to the raw gene expression matrix using scater in combination with SingleCellExperiment (version 1.28.1). We modeled the gene variance with scran (version 1.34.0) to identify the top 5,000 highly variable genes. We next merged all filtered samples and performed an additional sample-wide filtration step once again implementing the MAD approach to set a general threshold for minimum counts. The combined dataset contained 20,973 nuclei following quality filtration. For data integration across samples, we once again utilized the variational autoencoder from the scVI (**Construction of scRNA-seq atlas**).

###### snATAC-seq

ArchR (version 1.0.3) is an R package for processing and analyzing snATAC-seq data. Fragments files generated by cellranger-arc were imported, excluding mitochondrial chromosomes and retaining only barcodes corresponding to valid cells identified by cellranger-arc. Doublets were subsequently identified and removed using ArchR funtions addDoubletScores and filterDoublets. To account for differences in transcription start site (TSS) enrichment below a sample-specific threshold defined as the median TSS enrichment minus one standard deviation were excluded. We then applied a term frequency-inverse document frequency normalization using ArchR’s iterative latent semantic indexing. All filtered snATAC-seq samples were subsequently merged and Harmony (version 2.0.5) was applied to the dataset to mitigate batch effects.

##### Filtering of nuclei across datasets

Only nuclei with matched identities in both the snRNA-seq and snATAC-seq datasets were retained for downstream analyses of the corresponding modalities. The final matched snRNA-seq and snATAC-seq datasets comprised 14,683 nuclei of which 12,356 were malignant.

##### Annotation

###### snRNA-seq

Nuclei were clustered based on the scVI reduced dimensions via Leiden clustering producing 32 cell clusters and annotated as previously described (**Construction of scRNA-seq atlas**). At this stage, a low-quality nuclei cluster characterized by high mitochondrial read content was identified and removed from downstream analysis.

###### snATAC-seq

The Harmony-corrected dimensionality reduction was used for downstream clustering and visualization. Subcluster annotation was guided by the corresponding snRNA-seq data, with subcluster identities assigned to matched snATAC-seq nuclei based on their corresponding snRNA-seq annotations. Matched snRNA-seq data were integrated with the snATAC-seq data using addGeneExpressionMatrix. Mitochondrial genes were excluded prior to integration, and gene coordinates were provided as genomic ranges.

##### Peak calling

To identify regions of accessible chromatin, group coverages were generated using the ArchR addGroupCoverages package based on cell-state annotations, with a minimum of 20 nuclei per group (minCells) and ≥ 2 patient replicates (reproducibility). Reproducible chromatin accessibility peaks were subsequently identified using MACS3 (version 3.0.1) using addReproduciblePeakSet. Peaks were called using a q-value threshold of 0.1, with a shift of – 75 base pairs (bp) and an extension size of 150 bp. The MACS3 model and local lambda estimation were disabled, and peak summits were extended by 250 bp. Peaks detected in ≥ 2 replicates were retained to generate a reproducible peak set. A cell-by-peak accessibility matrix was generated for downstream analysis.

##### Inference of cis-regulatory elements

Putative cis-regulatory peak-to-gene associations were inferred using ArchR’s addPeak2GeneLinks. The Harmony-corrected dimensionality reduction was used to define neighbourhood structure and the integrated gene expression matrix used to calculate peak-gene correlations. Peak-to-gene associations were evaluated within a maximum genomic distance of 250 kbp, using 100 nearest neighbors and 500 knn interactions. An empirical p-value was incorporated to improve assessment of the significant of interchromosomal correlations. Peak-to-gene links were initially identified using a correlation threshold of 0.75 and an overlap cutoff of 0.8.

The resulting peak-to-gene links were subsequently retrieved using a Pearson correlation threshold of 0.45. To examine the cell-state distribution of linked regulatory elements, peak and gene expression matrices were aggregated into pseudobulk profiles and grouped into five k-means clusters based on their transcriptional patterns. GO enrichment analysis was performed for each regulatory cluster using goana from limma. All genes detected in the single-cell dataset were used as the background universe. Enriched Biological Process terms with p-value for over-representation < 0.05 and < 200 annotated genes were retained and summarized into semantic terms.

To independently validate regulatory interactions predicted by gene regulatory network inference, particularly the *NELL1*-HES4 regulatory link, motif annotations were inspected via pattern matching within called peaks linked to their nearest genes. N-box motif prevalence was calculated as the percentage of motif-positive *NELL1-*associated peaks.

##### Chromatin accessibility tracks

To enable visualization of chromatin accessibility in Integrated Genomics Viewer (version 3.0.0), BigWig files were generated using getGroupBW. Files were generated for cell types containing ≥ 20 nuclei, with read depth normalization based on reads in TSS regions. Gene loci of interest including *RIT2*+ tumor cell state marker *NELL1* were subsequently inspected in the viewer to visualize chromatin accessibility and assess regulatory features across cell populations.

##### Chromatin accessibility ternary plots

We evaluated relative promoter accessibility associated with the AC-like, OPC-like and NPC-like programs across the AC-like, OPC-like and *RIT2*+ tumor cell populations using ternary plots. For each cell nucleus in the snATAC-seq dataset, promoter peaks were annotated to their nearest gene and accessibility was binarized such that each gene was considered accessible if ≥ 1 promoter peak associated with that gene was detected. For each transcriptional program, the proportion of program genes with an accessible promoter was then calculated. Nuclei with no detectable promoter accessibility across any of the three programs were excluded and the resulting scores were normalized within each nucleus to sum to one. The relative promoter accessibility scores were visualized using ternary plots, colored according to tumor cell state using ggtern (version 4.0.0).

##### Chromosome 1p/19q analysis

Chromatin accessibility across chromosomes 1p and 19q was assessed by summarizing BigWig signal across 1Mbp genomic bins. Mean coverage was extracted for each track and normalized to the genome-wide median signal of the corresponding track. Accessibility across 1p and 19q was then compared between AC-like, OPC-like and *RIT2*+ tumor cell states.

#### Prediction algorithms

##### *RIT2*+ tumor cell prediction in ST

A penalized logistic regression model was trained using the tidymodels package (version 1.4.1) on malignant cell states. A 10-fold cross-validation was applied, with 25 % of malignant cells reserved as a testing set. To increase specificity, a prediction score cutoff of 0.70 was used to indicate *RIT2*+ tumor cells. Any malignant cell with a higher prediction score in the ST data was labelled as *RIT2*+ tumor cells. The identified niches of predicted *RIT2*+ tumor cells were then used to compare enrichment of *RIT2*+ tumor cells between different niches via an enrichment-based statistical test.

##### Immune adjustment approach, validation and application

To simulate the reduced gene panel of accompanying ST data, the reference single cell myeloid population was rerun through the consensus myeloid program calculator using only the genes within both the cNMF myeloid program signatures^37^ and the Xenium ST gene panels (n = 272). Combining this predicted myeloid program placement with the ground-truth myeloid program placement allows us to predict the displacement of individual cell predictions. Assuming that cells occupying a similar position of the program grid will be similarly displaced due to similarities in both their full and restricted gene profiles allows us to infer average adjustments needed to correct the displacement caused by the restricted gene panel. As evidenced by the general trends we observed in spatial displacement of predictions, cell displacements were reasonably consistent to enable this approach (**Figure 4D**). At a grid of points distributed every 5 units in both x and y directions, the average displacement was calculated for all cells in a surrounding 20 x 20 units square region. This was also performed separately for each sample, providing sample-specific grid adjustments. Adjustments of individual cells were calculated by interpolating between the surrounding adjustment values at the distributed points using interp package (version 1.1-6), without extrapolation enabled.

##### Immune adjustment of ST data

Raw immunomodulatory program scores for myeloid cells in the ST atlas were performed in the Shiny app, analogous to the restricted-panel snRNA-seq calculation. To estimate the displacement of the ST cells, the grid of adjustment values from the snRNA-seq data was repurposed. For cells from samples with matched snRNA-seq and ST data, sample-specific grid adjustment were used – otherwise, full-cohort grid adjustments were performed, with extrapolation enabled where necessary.

##### Immunomodulatory program spatial cooccurrence

To calculate the colocation of distinct immunomodulatory programs, ST cells were first characterized into distinct programs. Any cell with an absolute quadrant score greater than 20 was assigned to the corresponding progam (quadrant scores refers to the program grid before rotation, with the x axis corresponding to systemic inflammatory score minus complement immunosuppressive score, and the x axis corresponding to microglial inflammatory score minus scavenger immunosuppressive program). Due to the relative scarcity of scavenger immunosuppressive cells in the mIDH cohort, the threshold for this axis was reduced to 5. All other immune cells below these thresholds received no program label. A tiled window approach was used to describe cooccurrence of distinct programs. Raw counts of cells of each program were tallied across overlapping 1000 µm × 1000 µm tiles, with x- and y-plane shifts of 250 µm between overlapping tiles. Relative pairwise proportions of programs and overall program counts were then collected.

##### T-ROS niche prediction, validation and application

A penalized logistic regression was performed on a downsampled subset of OPC-like and AC-like malignant cells within the ST cohort using the tidymodels package (version 1.4.1). Downsampling followed several principles: for deeply profiled samples with multiple z-layers, only cells from the 1st z-layer were included; All cells belonging in the T-ROS niche were included; among cells from other niches, a maximum of 45,000 cells per niche were included, split equally across samples by limiting sample thresholds to 2,000 or 4,000 cells depending on overall niche abundance. Additionally, samples A3 and A6 were held out of training completely as representative positive and negative controls, respectively. Of the 434,401 downsampled cells, 75 % (325,800) underwent 10-fold cross validation, with the remaining 25 % (107,601) used as a testing set. 50 penalty values evenly spaced along a log-scale of 10^−5^ to 10^5^ were assessed, with minimal penalization conferring optimal performance. Hence, a penalty of 0.00001 was chosen for the resulting regression model.

