## Supplementary Figures for "Spatial metabolic ecosystems shape the architecture of *IDH*-mutant glioma"

#### Supplementary Figures and Figure Legends

Figure S1

Figure S2

Figure S3

Figure S4

Figure S5

Figure S6

### Figure S1

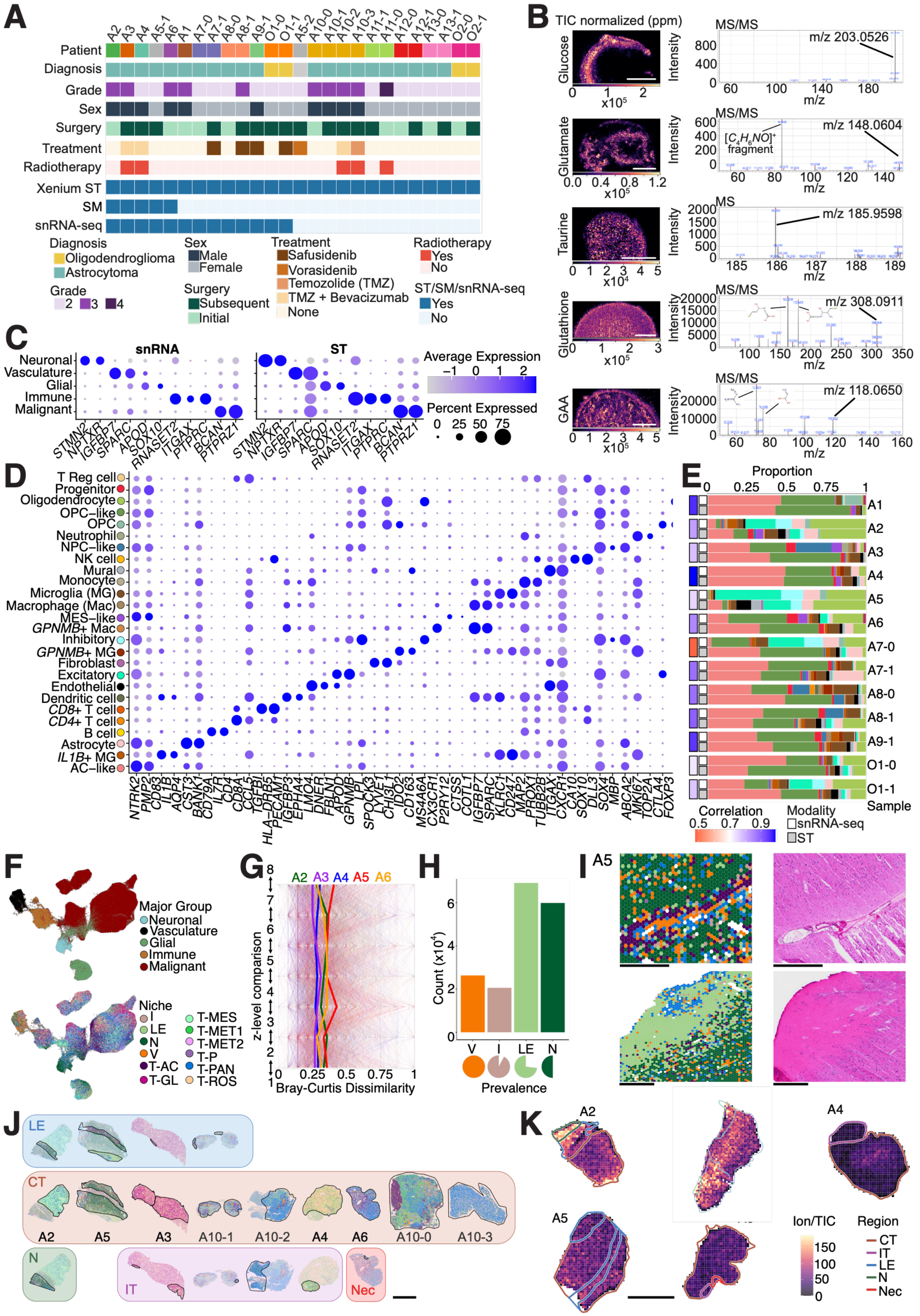

A
Patient
Diagnosis
Grade
Sex
Surgery
Treatment
Radiotherapy
Xenium ST
SM
snRNA-seq
Diagnosis
Oligodendroglioma
Astrocytoma
Sex
Male
Female
Treatment
Safusidenib
Vorasidenib
Temozolide (TMZ)
TMZ + Bevacizumab
None
Radiotherapy
Yes
No
Grade
2 3 4
Surgery
Subsequent
Initial
ST/SM/snRNA-seq
Yes
No
B
TIC normalized (ppm)
Glucose
Glutamate
Taurine
Glutathione
GAA
MS/MS
MS
m/z 203.0526
m/z 148.0604
m/z 185.9598
m/z 308.0911
m/z 118.0650
Intensity
m/z
C
snRNA
ST
Neuronal
Vasculature
Glial
Immune
Malignant
Average Expression
Percent Expressed
D
E
Proportion
Correlation
Modality
snRNA-seq
ST
Sample
F
Major Group
Neuronal
Vasculature
Glial
Immune
Malignant
Niche
G
z-level comparison
Bray-Curtis Dissimilarity
H
Count (x10⁴)
Prevalence
I
J
LE
CT
N
IT
Nec
K
Ion/TIC
Region
CT
IT
LE
N
Nec

**Figure S1. related to Figure 1**

**A.** Patient metadata of samples profiled with ST, including aspects of diagnosis, surgical timepoint and treatment, as well as multi-omic profiling strategy. **B.** Metabolite standards analyzed by MS/MS or MS (Taurine only). Left, image of spotted metabolite with TIC normalized intensity. Scale, 500 µm. Right, Mass spectra trace with dominant peak highlighted confirming metabolite peak or fragment identify. **C.** Dot plot of log-normalized z-scores of gene expression for markers of major cell groups, both in snRNA-seq data (left) and ST data (right). **D.** Dot plot of log-normalized z-scores of gene expression for markers of malignant and brain-resident cell types in ST data. **E.** Bar plot of cell type proportions for samples with matched ST and snRNAseq data. Correlation of proportions of shared cell type annotations is calculated for each sample pair. **F.** UMAP of ST cohort, colored by major cell group (above) and GraphSAGE niche (below). **G.** Bray-Curtis dissimilarity of niche composition of aligned 1000 µm by 1000 µm regions of adjacent sections from deeply profiled samples (n = 5). Median patient scores in bold. **H.** Abundance and prevalence of non-malignant associated niches. **I.** Left, representative ST LE regions of sample A5, colored by niche. Right, adjacent H&E sections of sampled tissue regions. Scale, 1 mm. **J.** ST samples split by annotated histopathological regions, colored by niche distribution. Scale, 5 mm. **K.** TIC-normalized abundance of 2-HG (m/z value: 174.998) across all 5 deeply profiled samples, with histopathological regions overlaid. Scale, 5 mm.

### Figure S2

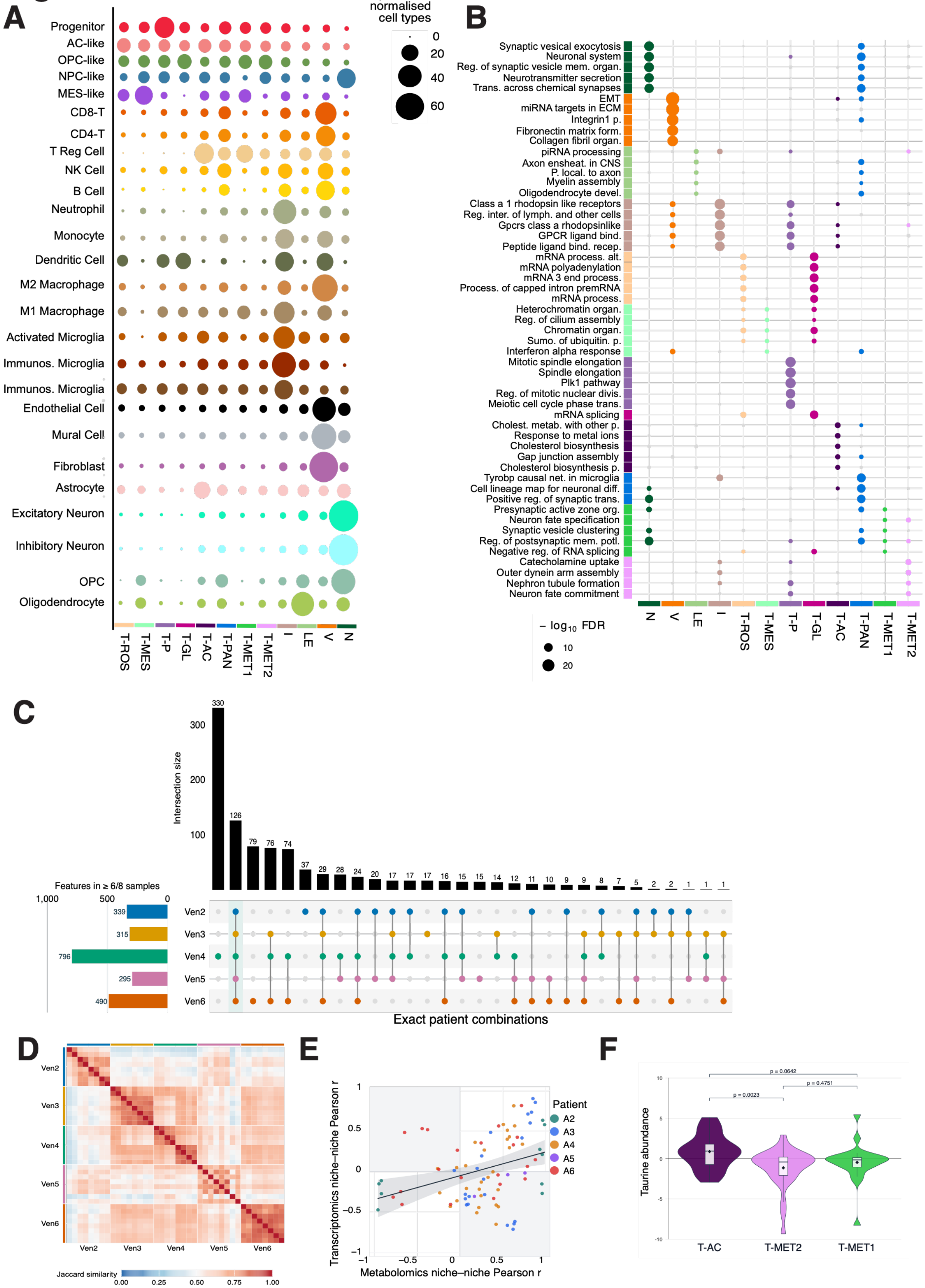
A
B
C
D
E
F
Proportion of normalised cell types
Progenitor
AC-like
OPC-like
NPC-like
MES-like
CD8-T
CD4-T
T Reg Cell
NK Cell
B Cell
Neutrophil
Monocyte
Dendritic Cell
M2 Macrophage
M1 Macrophage
Activated Microglia
Immunos. Microglia
Endothelial Cell
Mural Cell
Fibroblast
Astrocyte
Excitatory Neuron
Inhibitory Neuron
OPC
Oligodendrocyte
T-ROS T-MES T-P T-GL T-AC T-PAN T-MET1 T-MET2 I LE V N
Synaptic vesical exocytosis
Neuronal system
Reg. of synaptic vesicle mem. organ.
Neurotransmitter secretion
Trans. across chemical synapses
EMT
miRNA targets in ECM
Integrin1 p.
Fibronectin matrix form.
Collagen fibril organ.
piRNA processing
Axon ensheat. in CNS
P. local. to axon
Myelin assembly
Oligodendrocyte devel.
Class a 1 rhodopsin like receptors
Reg. inter. of lymph. and other cells
Gpcrs class a rhodopsinlike
GPCR ligand bind.
Peptide ligand bind. recep.
mRNA process. alt.
mRNA polyadenylation
mRNA 3 end process.
Process. of capped intron premRNA
mRNA process.
Heterochromatin organ.
Reg. of cilium assembly
Chromatin organ.
Sumo. of ubiquitin. p.
Interferon alpha response
Mitotic spindle elongation
Spindle elongation
Plk1 pathway
Reg. of mitotic nuclear divis.
Meiotic cell cycle phase trans.
mRNA splicing
Cholest. metab. with other p.
Response to metal ions
Cholesterol biosynthesis
Gap junction assembly
Cholesterol biosynthesis p.
Tyrobp causal net. in microglia
Cell lineage map for neuronal diff.
Positive reg. of synaptic trans.
Presynaptic active zone org.
Neuron fate specification
Synaptic vesicle clustering
Reg. of postsynaptic mem. potl.
Negative reg. of RNA splicing
Catecholamine uptake
Outer dynein arm assembly
Nephron tubule formation
Neuron fate commitment
− log10 FDR
N V LE I T-ROS T-MES T-P T-GL T-AC T-PAN T-MET1 T-MET2
Intersection size
Features in ≥ 6/8 samples
Ven2 Ven3 Ven4 Ven5 Ven6
Exact patient combinations
Jaccard similarity
Transcriptomics niche–niche Pearson r
Metabolomics niche–niche Pearson r
Patient
A2 A3 A4 A5 A6
Taurine abundance
p = 0.0642
p = 0.0023
p = 0.4751
T-AC T-MET2 T-MET1

**Figure S2. related to Figure 2**

**A.** Enrichment plot of cell type annotations in each GraphSAGE niche. **B.** Top upregulated pathways per spatial niche. Heatmap of the union of the five most significantly upregulated gene sets from each niche, showing -Log10FDR from cell-type-adjusted cameraPR tests of each niche against the mean of the other 11 (BH-corrected across gene sets within each niche; *FDR < 0.05). **C.** Jaccard similarity of metabolite profiles of different samples and their 8 different profiled layers. **D.** Upset plot of m/z peaks identified across all 8 layers of multiple patient samples. **E.** Correlation of Pearson's R values for SM niche-niche pairs and ST niche-niche pairs, from (Figure 2C). **F.** Taurine abundance in T-MET1 compared to T-MET2 niche.

### Figure S3

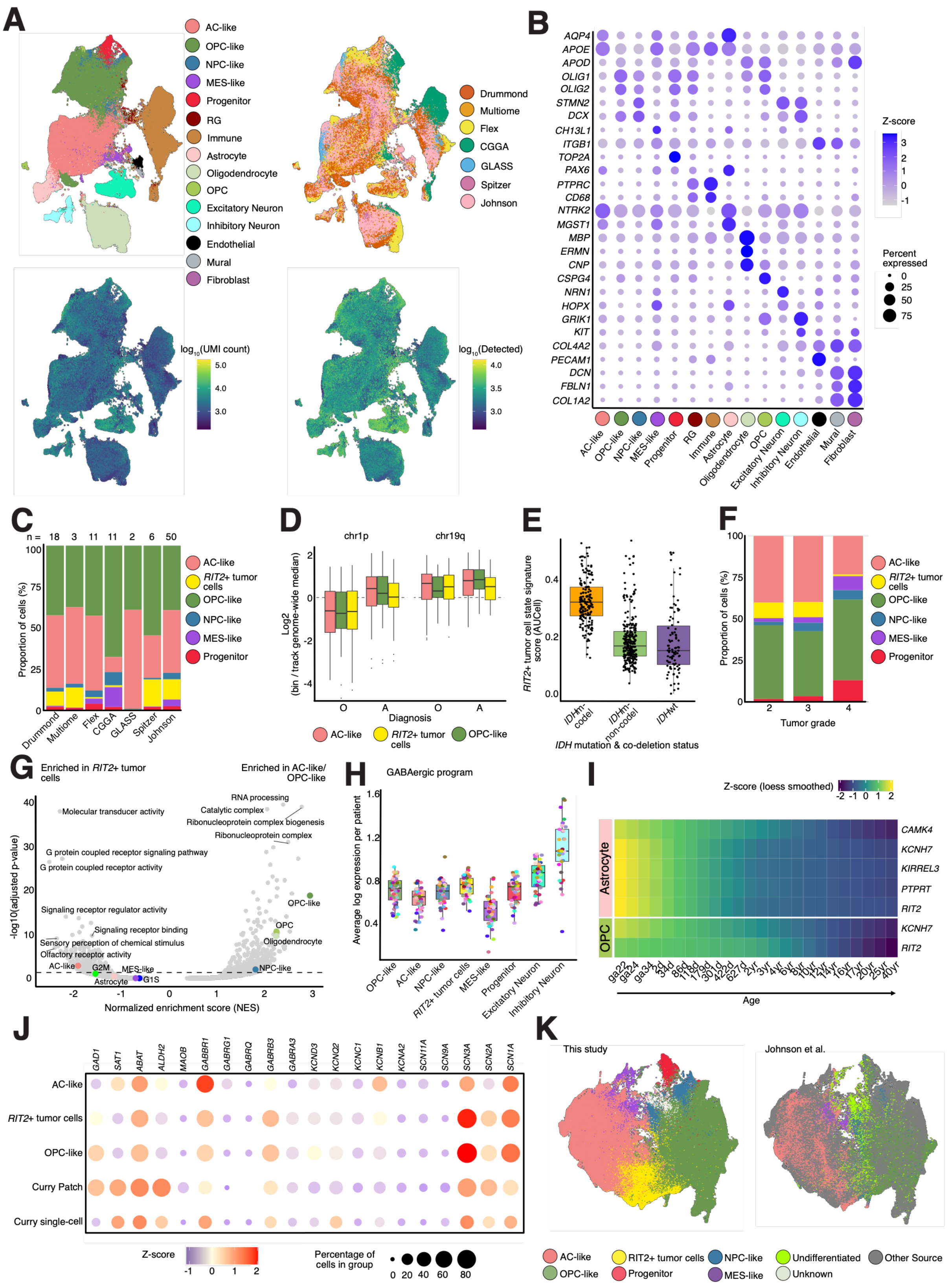
A
AC-like
OPC-like
NPC-like
MES-like
Progenitor
RG
Immune
Astrocyte
Oligodendrocyte
OPC
Excitatory Neuron
Inhibitory Neuron
Endothelial
Mural
Fibroblast
Drummond
Multiome
Flex
CGGA
GLASS
Spitzer
Johnson
log10(UMI count)
5.0
4.0
3.0
log10(Detected)
4.0
3.5
3.0
B
AQP4
APOE
APOD
OLIG1
OLIG2
STMN2
DCX
CH13L1
ITGB1
TOP2A
PAX6
PTPRC
CD68
NTRK2
MGST1
MBP
ERMN
CNP
CSPG4
NRN1
HOPX
GRIK1
KIT
COL4A2
PECAM1
DCN
FBLN1
COL1A2
Z-score
3
2
1
0
-1
Percent expressed
0
25
50
75
C
n = 18 3 11 11 2 6 50
Proportion of cells (%)
100
75
50
25
0
Drummond
Multiome
Flex
CGGA
GLASS
Spitzer
Johnson
AC-like
RIT2+ tumor cells
OPC-like
NPC-like
MES-like
Progenitor
D
chr1p
chr19q
Log2 (bin / track genome-wide median)
Diagnosis
O
A
E
RIT2+ tumor cell state signature score (AUCell)
IDHm-codel
IDHm-non-codel
IDHwt
IDH mutation & co-deletion status
F
Proportion of cells (%)
Tumor grade
2
3
4
G
Enriched in RIT2+ tumor cells
Enriched in AC-like/ OPC-like
-log10(adjusted p-value)
Normalized enrichment score (NES)
Molecular transducer activity
RNA processing
Catalytic complex
Ribonucleoprotein complex biogenesis
Ribonucleoprotein complex
G protein coupled receptor signaling pathway
G protein coupled receptor activity
Signaling receptor regulator activity
Signaling receptor binding
Sensory perception of chemical stimulus
Olfactory receptor activity
AC-like
G2M
MES-like
Astrocyte
G1S
NPC-like
OPC
Oligodendrocyte
OPC-like
H
GABAergic program
Average log expression per patient
OPC-like
AC-like
NPC-like
RIT2+ tumor cells
MES-like
Progenitor
Excitatory Neuron
Inhibitory Neuron
I
Z-score (loess smoothed)
Astrocyte
OPC
CAMK4
KCNH7
KIRREL3
PTPRT
RIT2
KCNH7
RIT2
ga22
ga24
ga34
2d
34d
86d
118d
179d
301d
422d
627d
2yr
3yr
4yr
6yr
8yr
10yr
12yr
14yr
16yr
17yr
20yr
25yr
40yr
Age
J
GAD1
SAT1
ABAT
ALDH2
MAOB
GABBR1
GABRG1
GABRQ
GABRB3
GABRA3
KCND3
KCNQ2
KCNC1
KCNB1
KCNA2
SCN11A
SCN9A
SCN3A
SCN2A
SCN1A
AC-like
RIT2+ tumor cells
OPC-like
Curry Patch
Curry single-cell
Z-score
-1 0 1 2
Percentage of cells in group
0 20 40 60 80
K
This study
Johnson et al.
AC-like
RIT2+ tumor cells
NPC-like
Undifferentiated
Other Source
OPC-like
Progenitor
MES-like
Unknown

**Figure S3. related to Figure 3**

**A.** UMAP plot depicting grade 2 and 3 mIDH glioma reference atlas of cells and nuclei spanning malignant and non-malignant states. Top left: annotated UMAP clusters of glial tumor and non-malignant cell states; top right: dataset source; bottom left: UMAP colored by UMI count ($\log_{10}$ transformed); bottom right: UMAP colored by number of detected genes ($\log_{10}$ transformed). **B.** Dot plot of z-score of log-normalized gene expression of well-established markers across different cell types. **C.** Proportion of glial tumor cell states across dataset source. **D.** Log-normalized chromatin accessibility across chromosomes 1p and 19q compared between AC-like, OPC-like and *RIT2*+ tumor cell populations using 1 Mbp genomic bins. **E.** Gene set enrichment scores (AUCell) of the *RIT2*+ tumor cell state signature across tumors of distinct *IDH* mutation and chromosome 1p and 19q co-deletion status from the TCGA bulk RNA sequencing low grade glioma cohort. **F.** Proportion of glial tumor cell states across mIDH glioma tumor grades. **G.** Comparison of AC-like and OPC-like to *RIT2*+ tumor cells in the reference atlas using GSEA via ranked lists according to DEG results. Each dot represents a gene-set and color indicates whether the gene-set belongs to a glioma program and top enriched pathways are annotated. X axis shows the GSEA normalized enrichment score (NES), Y axis shows the adjusted p-value. Line indicates significance threshold (adjusted p-value 0.05). **H.** Box plot of GABAergic average log normalized gene expression per patient in different cell populations of mIDH reference atlas. **I.** Heatmap of LOESS smoothed average $\log_2$ expression of candidate *RIT2*+ tumor cell state markers across developmental stages of normal Astrocyte and OPC populations in the scRNA- seq cohort from *Herring* et al. **J.** Dot plot of z-score average log-normalized expression of key markers described by *Curry* et al., for an OPC-GABA population from Patch-sequencing and scRNA- seq compared to AC-like, OPC-like and *RIT2*+ tumor cells from the mIDH reference atlas. **K.** UMAP plots depicting comparison of mIDH reference atlas to 'Undifferentiated' population described by *Johnson* et al. Left: assigned glioma cell state annotation from this study; right: assigned glioma cell state annotations by *Johnson* et al. Cells/nuclei integrated from other dataset sources (Other Source) labeled.

### Figure S4

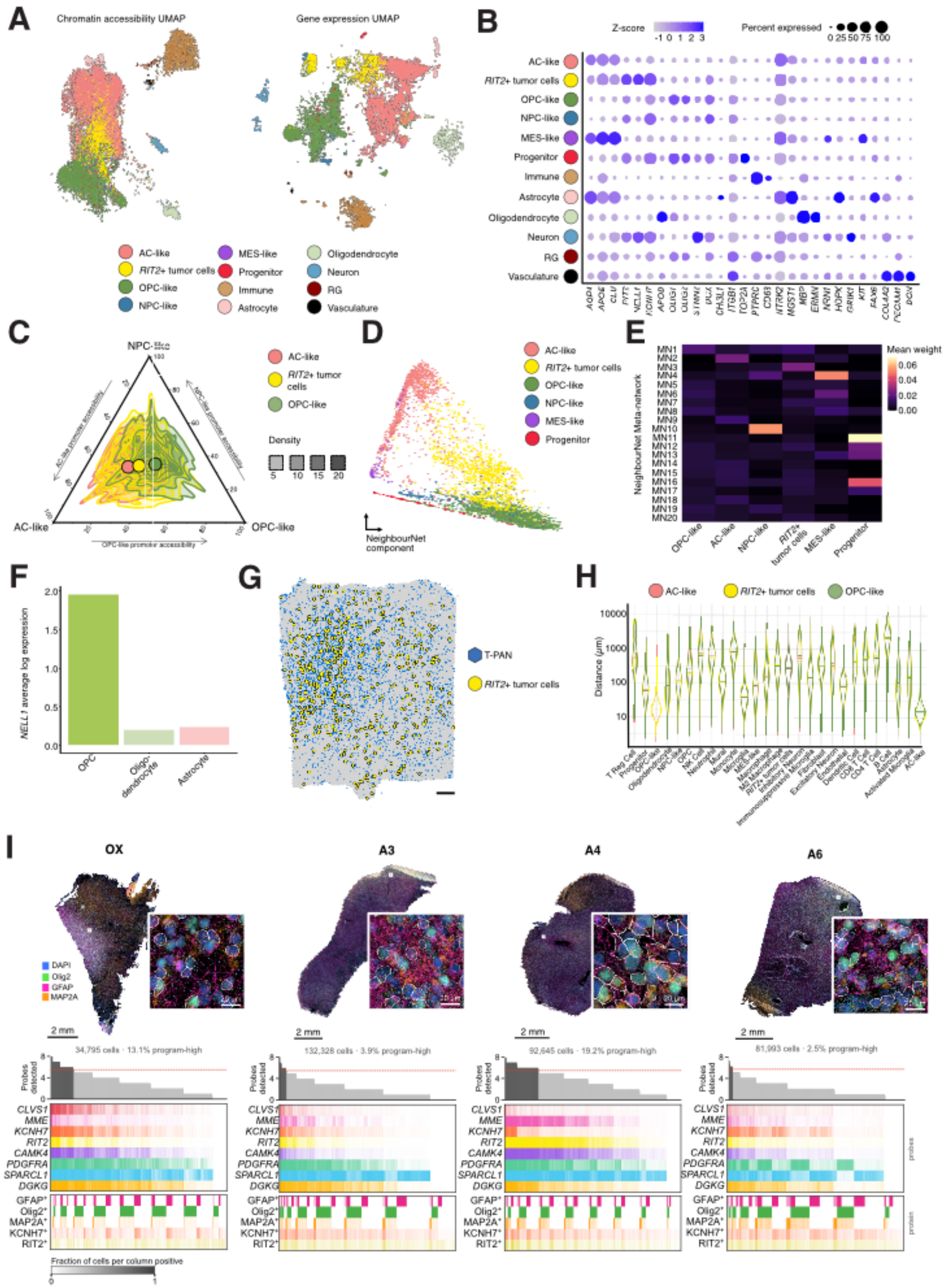
A
Chromatin accessibility UMAP
Gene expression UMAP
AC-like
RIT2+ tumor cells
OPC-like
NPC-like
MES-like
Progenitor
Immune
Astrocyte
Oligodendrocyte
Neuron
RG
Vasculature
B
Z-score
Percent expressed
C
NPC-like
AC-like
OPC-like
Density
D
NeighbourNet component
E
NeighbourNet Meta-network
Mean weight
F
NELL1 average log expression
OPC
Oligo-dendrocyte
Astrocyte
G
T-PAN
RIT2+ tumor cells
H
Distance (μm)
I
OX
A3
A4
A6
DAPI
Olig2
GFAP
MAP2A
2 mm
34,795 cells · 13.1% program-high
132,328 cells · 3.9% program-high
92,645 cells · 19.2% program-high
81,993 cells · 2.5% program-high
Probes detected
CLVS1
MME
KCNH7
RIT2
CAMK4
PDGFRA
SPARCL1
DGKG
GFAP+
Olig2+
MAP2A+
KCNH7+
RIT2+
Fraction of cells per column positive

**Figure S4. related to Figure 3**

**A.** UMAP plots depicting ATAC-seq (chromatin accessibility) and snRNA-seq (gene expression) Multiome data included in mIDH reference atlas. Left: annotated chromatin accessibility UMAP clusters; right: annotated gene expression UMAP clusters from the same nuclei. **B.** Dot plot of z-score of log-normalized expression of well-established and candidate *RIT2*+ tumor cell state markers across different cell types from the Multiome. **C.** Ternary plot depicting promoter chromatin accessibility of glioma programs; AC-like, OPC-like and NPC-like, in AC-like, *RIT2*+ tumor cells and OPC-like tumor cells. **D.** NeighbourNet inferred components space colored by downsampled glioma cell states from oligodendroglioma tumors of the mIDH reference atlas. **E.** NeighbourNet mean weight of inferred meta-networks for each glioma cell state. **F.** Average log-normalized expression of *NELL1* across OPC, oligodendrocyte and astrocyte populations from the single-cell RNA sequencing cohort from *Herring* et al. **G.** Spatial transcriptomics plot depicting co-localization of *RIT2*+ tumor cells to the T-PAN GraphSAGE niche in patient O2-1. Scale, 1 mm. **H.** Distance (in µm) of cell populations from the mIDH reference atlas to OPC-like, AC-like and *RIT2*+ tumor cells. **I.** *RIT2*+ tumor cell RNA and protein co-detection. Top: Whole-section overviews of the patient OX, A3, A4 and A6 after RNAscope sequential IF (DAPI, GFAP, Olig2, MAP2A). Scale bars, 2 mm. Insets, 20 µm. A field from each section (white box on the section) whole-cell segmentation (grey outlines) and individual RNAscope transcripts of the 8 program probes. Cells detecting at least 6 of the 8 probes (program-high) are outlined in white. Scale bars, 20 µm. Bottom: Co-detection matrices with cells sorted by the number of probes detected and grouped into 400 consecutive columns (87-331 cells per column). First row: Mean number of probes detected per column; black, program-high columns; dashed line, program-high threshold. Second row: Fraction of cells in the column detecting the probe (at least 2 estimated transcripts per cell; white, 0; full colour, 1). Bottom row: Fraction of cells positive for the IF markers GFAP,

Olig2 and MAP2A and for the program-gene proteins KCNH7 and RIT2. Percentage of program-high cells per section is given above each matrix.

### Figure S5

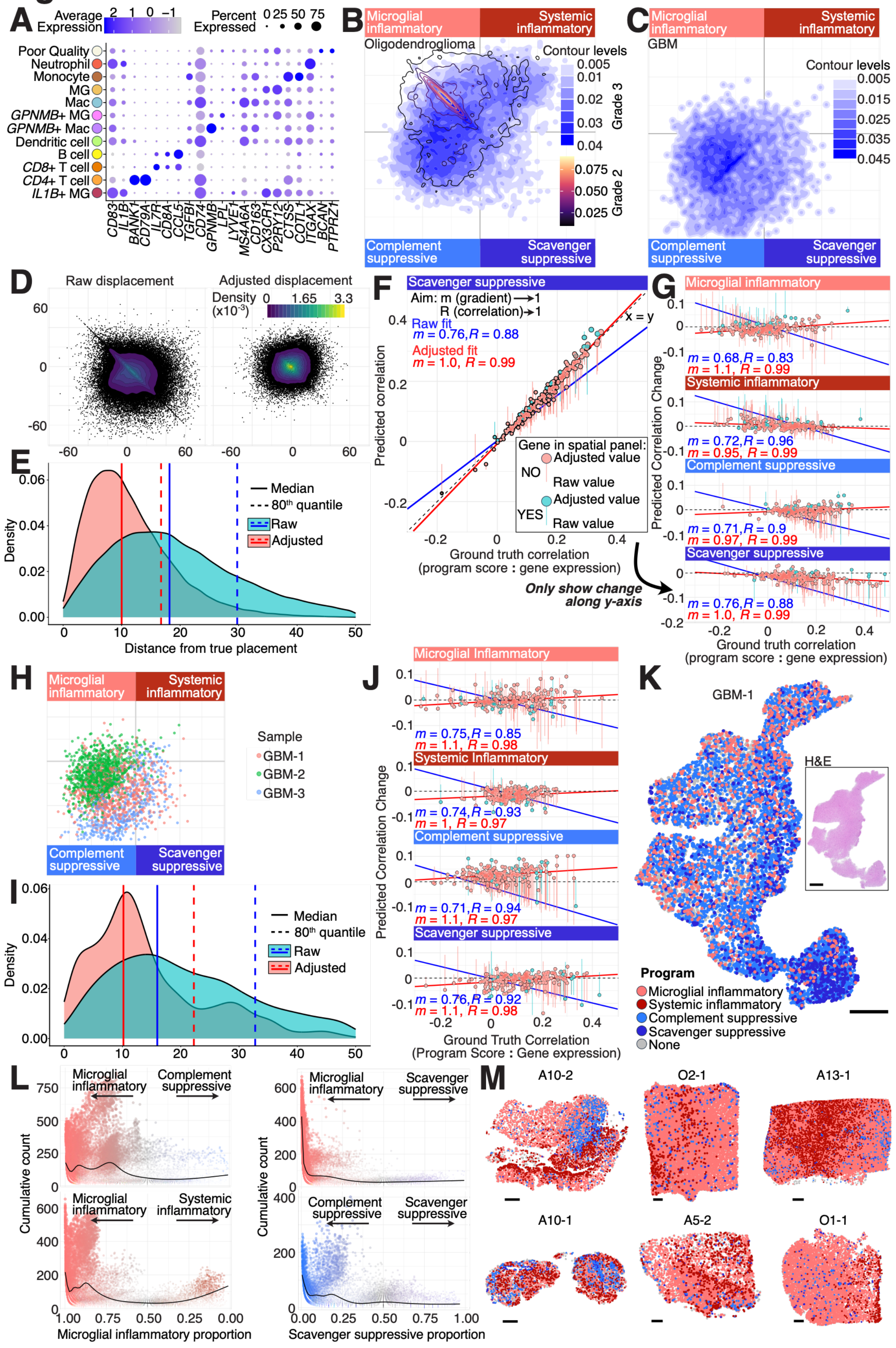

A
Average Expression
2 1 0 −1
Percent Expressed
0 25 50 75
Poor Quality
Neutrophil
Monocyte
MG
Mac
GPNMB+ MG
GPNMB+ Mac
Dendritic cell
B cell
CD8+ T cell
CD4+ T cell
IL1B+ MG
CD83 IL1B BANK1 CD79A IL7R CD8A CCL5 TGFBI CD74 GPNMB LPL LYVE1 MS4A6A CD163 CX3CR1 P2RY12 CTSS COTL1 ITGAX BCAN PTPRZ1
B
Microglial inflammatory
Systemic inflammatory
Oligodendroglioma
Contour levels
0.005 0.01 0.02 0.03 0.04
Grade 3
0.075 0.050 0.025
Grade 2
Complement suppressive
Scavenger suppressive
C
Microglial inflammatory
Systemic inflammatory
GBM
Contour levels
0.005 0.015 0.025 0.035 0.045
Complement suppressive
Scavenger suppressive
D
Raw displacement
Adjusted displacement
Density (x10^-3)
0 1.65 3.3
60 0 -60
-60 0 60 -60 0 60
E
Density
Median
80th quantile
Raw
Adjusted
Distance from true placement
F
Scavenger suppressive
Aim: m (gradient)→1
R (correlation)→1
x = y
Raw fit
m = 0.76, R = 0.88
Adjusted fit
m = 1.0, R = 0.99
Gene in spatial panel:
NO
Adjusted value
Raw value
YES
Adjusted value
Raw value
Predicted correlation
Ground truth correlation
(program score : gene expression)
Only show change along y-axis
G
Microglial inflammatory
m = 0.68, R = 0.83
m = 1.1, R = 0.99
Systemic inflammatory
m = 0.72, R = 0.96
m = 0.95, R = 0.99
Complement suppressive
m = 0.71, R = 0.9
m = 0.97, R = 0.99
Scavenger suppressive
m = 0.76, R = 0.88
m = 1.0, R = 0.99
Predicted Correlation Change
Ground truth correlation
(program score : gene expression)
H
Microglial inflammatory
Systemic inflammatory
Sample
GBM-1
GBM-2
GBM-3
Complement suppressive
Scavenger suppressive
I
Density
Median
80th quantile
Raw
Adjusted
J
Microglial Inflammatory
m = 0.75, R = 0.85
m = 1.1, R = 0.98
Systemic Inflammatory
m = 0.74, R = 0.93
m = 1, R = 0.97
Complement suppressive
m = 0.71, R = 0.94
m = 1.1, R = 0.97
Scavenger suppressive
m = 0.76, R = 0.92
m = 1.1, R = 0.98
Predicted Correlation Change
Ground Truth Correlation
(Program Score : Gene Expression)
K
GBM-1
H&E
Program
Microglial inflammatory
Systemic inflammatory
Complement suppressive
Scavenger suppressive
None
L
Microglial inflammatory
Complement suppressive
Scavenger suppressive
Systemic inflammatory
Complement suppressive
Scavenger suppressive
Cumulative count
Microglial inflammatory proportion
Scavenger suppressive proportion
M
A10-2
O2-1
A13-1
A10-1
A5-2
O1-1

**Figure S5. related to Figure 4**

**A.** Dot plot of z-score of average log-normalized expression of immune cell type markers in snRNA-seq dataset. **B.** 2D density of immunomodulatory program distribution of snRNA-seq oligodendroglioma samples, split into grade 3 (blue gradient) and grade 2 ('magma' gradient) samples. **C.** 2D density of immunomodulatory program distribution of snRNA-seq GBM samples. **D.** Scatterplot of displacement of snRNA-seq myeloid cells in immunomodulatory program space when predicting placement with the restricted gene panel (raw, left) and after adjustment using calibration described in **Methods** (adjusted, right). **E.** Density of distance from true placement in immunomodulatory program space for snRNA-seq myeloid cells, using either unadjusted positions from the restricted ST gene panel (raw, blue) or adjusted position using the calibration method (adjusted, red). **F.** Example assessment of adjustment correction for the scavenger immunosuppressive program. Scatterplot comparing ground-truth correlations with the scavenger immunosuppressive program score against correlations predicted from raw and adjusted restricted-panel data for the 200 most strongly correlated genes. For each gene, the dot indicates the adjusted prediction, while the unmarked end of the connecting vertical line indicates the raw prediction. Genes are colored according to their inclusion in the restricted ST panel (light blue, included; light red, excluded). Linear regression lines are fitted to the raw (blue) and adjusted (red) predictions, with the corresponding slopes (m) and correlation coefficients (R) indicated. **G.** Assessment of adjustment correction for all four immunomodulatory programs, analogous to (F). Ground truth program score correlation of the top 200 genes for each program is compared to the predicted change in correlation, from raw values to adjusted values. Gradient and correlation values are calculated as described in (F). **H.** Immunomodulatory program distribution of snRNA-seq GBM samples, colored by sample. **I.** Density of distance from true placement in immunomodulatory program space for snRNA-seq GBM myeloid cells, using either unadjusted positions from the restricted ST gene panel (raw, blue) or adjusted position using the calibration method (adjusted, red). **J.** Assessment of adjustment correction for all four immunomodulatory programs in GBM data as described in (G). Ground truth program score correlation of the top 200 genes for each program is compared to the predicted change in correlation, from raw values to adjusted values. Gradient and correlation values are calculated as described in (F). **K.** Spatial distribution of immunomodulatory programs in GBM-1. Myeloid cells colored by program if program score is ≥ 20 (or ≥ 5 for scavenger immunosuppressive cells). Inset, H&E staining of GBM-1. Scale bars, 1mm. **L.** Comparison of overall number of ST myeloid cells within program pairs, and the proportion of each program within these pairs. Only ST myeloid cells with program scores of ≥ 20 were included in analysis, or ≥ 5 for scavenger immunosuppressive cells. Measured regions were 1000 µm by 1000 µm across all ST samples. **M.** Spatial distribution of myeloid cells colored by program across 6 patient samples. Scale bars, 1mm.

### Figure S6

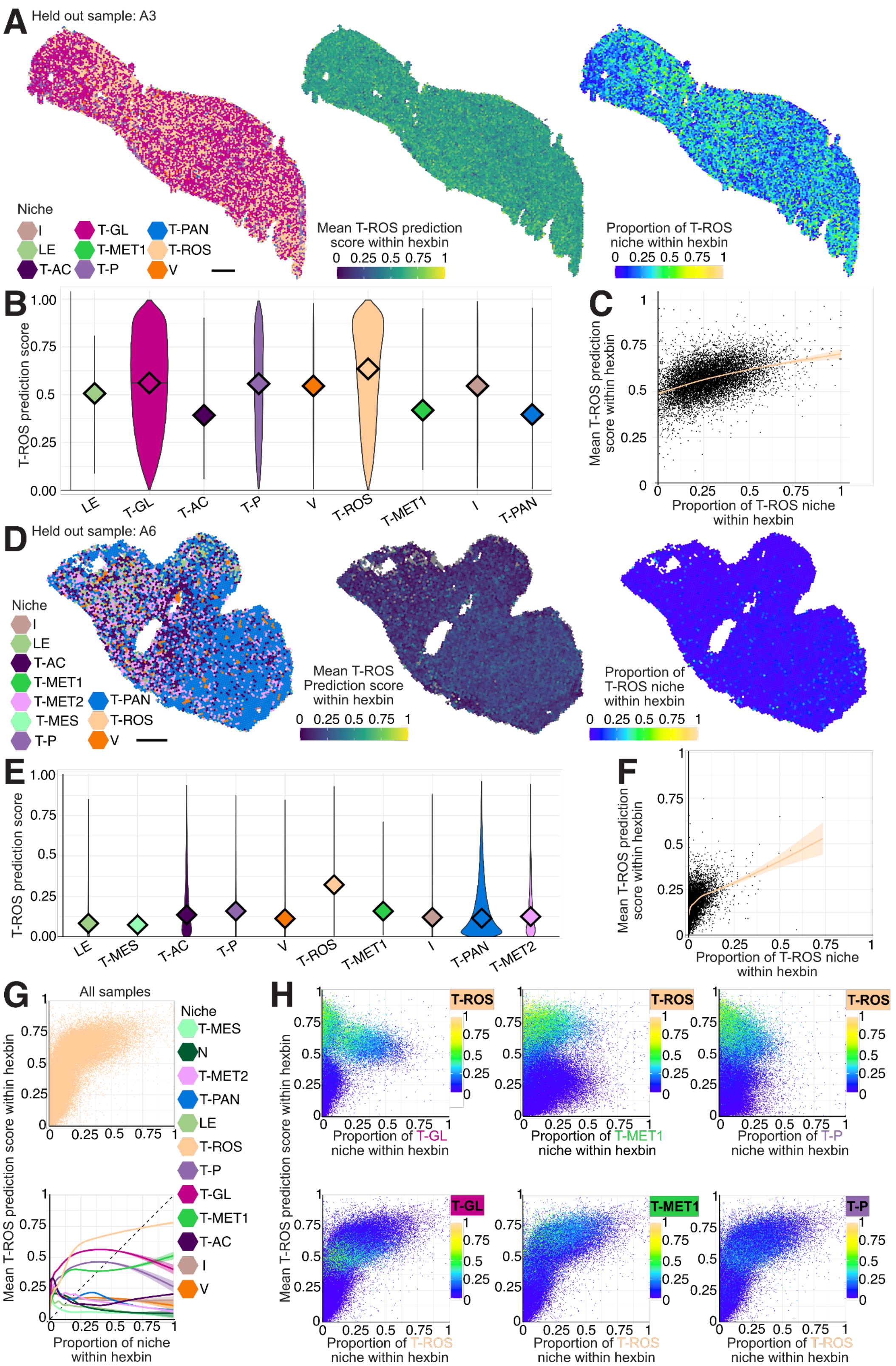
A
Held out sample: A3
Niche
I
LE
T-AC
T-GL
T-MET1
T-P
T-PAN
T-ROS
V
Mean T-ROS prediction score within hexbin
0 0.25 0.5 0.75 1
Proportion of T-ROS niche within hexbin
0 0.25 0.5 0.75 1
B
T-ROS prediction score
1.00 0.75 0.5 0.25 0.00
LE T-GL T-AC T-P V T-ROS T-MET1 I T-PAN
C
Mean T-ROS prediction score within hexbin
Proportion of T-ROS niche within hexbin
D
Held out sample: A6
Niche
I
LE
T-AC
T-MET1
T-MET2
T-MES
T-P
T-PAN
T-ROS
V
Mean T-ROS Prediction score within hexbin
0 0.25 0.5 0.75 1
Proportion of T-ROS niche within hexbin
0 0.25 0.5 0.75 1
E
T-ROS prediction score
1.00 0.75 0.5 0.25 0.00
LE T-MES T-AC T-P V T-ROS T-MET1 I T-PAN T-MET2
F
Mean T-ROS prediction score within hexbin
Proportion of T-ROS niche within hexbin
G
All samples
Mean T-ROS prediction score within hexbin
Proportion of niche within hexbin
Niche
T-MES
N
T-MET2
T-PAN
LE
T-ROS
T-P
T-GL
T-MET1
T-AC
I
V
H
Mean T-ROS prediction score within hexbin
T-ROS
Proportion of T-GL niche within hexbin
Proportion of T-MET1 niche within hexbin
Proportion of T-P niche within hexbin
T-GL
T-MET1
T-P
Proportion of T-ROS niche within hexbin

**Figure S6. related to Figure 5**

**A.** From left to right: GraphSAGE niche distribution of held-out patient A3; distribution of T-ROS logistic regression prediction score across hexbins of patient A3, calculated as the mean T-ROS prediction score across all OPC-like and AC-like cells within each hexbin (hexbins without any OPC-like or AC-like tumor cells are colored grey); distribution of true T-ROS proportion within each hexbin, calculated as the proportion of node labels within the hexbin labelled as T-ROS. Scale, 1mm. **B.** Violin plot of predicted T-ROS signature scores for AC-like and OPC-like cells in patient A3, split by assigned GraphSAGE niche. Median indicated by diamond. **C.** Scatterplot of hexbins in patient A3 comparing proportion of T-ROS niche within hexbin nodes and mean T-ROS prediction score within the hexbin, with a fitted generalized linear model overlaid. **D.** From left to right: GraphSAGE niche distribution of held-out patient A6; distribution of T-ROS logistic regression prediction score across hexbins of patient A6, calculated as the mean T-ROS prediction score across all OPC-like and AC-like cells within each hexbin (hexbins without any OPC-like or AC-like tumor cells are colored grey); distribution of true T-ROS proportion within each hexbin, calculated as the proportion of node labels within the hexbin labelled as T-ROS. Scale, 1mm. **E.** Violin plot of predicted T-ROS signature scores for AC-like and OPC-like cells in patient A6, split by assigned GraphSAGE niche. Median indicated by diamond. **F.** Scatterplot of hexbins in patient A6 comparing the proportion of T-ROS niche within hexbin nodes and mean T-ROS prediction score within the hexbin, with a fitted generalized linear model overlaid. **G.** Above, scatterplot of hexbins across all samples comparing the proportion of T-ROS niche within hexbin nodes and mean T-ROS prediction score within the hexbin. Below, fitted generalized linear models comparing the proportion of each GraphSAGE niche within hexbin nodes and mean T-ROS prediction score within the hexbin. **H.** Top row: scatterplots of hexbins across all samples comparing the proportion of 3 niches (T-GL, T-MET1, and T-P, respectively) within a hexbin, and its mean T-ROS prediction score – colored by the proportion of T-ROS niche nodes within the hexbin. Bottom row: scatterplots of hexbins across all samples comparing the proportion of T-ROS niche nodes within the hexbin, and its mean T-ROS prediction score – colored by the proportion of 3 niches (T-GL, T-MET1, and T-P, respectively) within the hexbin.
